# NR4A3 deficiency in CD8^+^ T cells improves adoptive T cell therapy of cancer

**DOI:** 10.1101/2023.04.21.537841

**Authors:** Livia Odagiu, Sarah Mezrag, Dave Maurice De Sousa, Salix Boulet, Nathalie Labrecque

## Abstract

NR4A3 is a transcription factor that is rapidly induced in CD8^+^ T cells following antigenic recognition. We have previously shown that NR4A3 deficiency induces an early molecular program that promotes memory generation and enhances effector functions, which are two essential attributes for the success of adoptive cell therapy (ACT). Therefore, we tested the hypothesis that *Nr4a3*^-/-^ CD8^+^ T cells would have outstanding efficacy in ACT of cancer. Our results show that ACT of melanoma-bearing mice with *Nr4a3*^-/-^ effector CD8^+^ T cells provides a better tumor control than their wild-type counterpart. The therapeutic effect observed with *Nr4a3*^-/-^ effector CD8^+^ T cells is even better than the one observed with ACT of *Nr4a3*^+/+^ effector CD8^+^ T cells in combination with anti-PD-L1 treatment. scRNA-seq analysis reveals a huge heterogeneity of tumor-infiltrating lymphocytes (TILs) states following ACT. The better tumor control observed with ACT of *Nr4a3*^-/-^ CD8^+^ effectors without anti-PD-L1 treatment correlates with an enrichment of TILs within the clusters that are associated with the anti-PD-L1 response of wild-type TILs. Moreover, the clusters that are enriched in *Nr4a3*^-/-^ TILs are the ones that are enriched for effector functions. Furthermore, *Nr4a3*^-/-^ and *Nr4a3*^+/+^ effectors generate distinct progenitor populations. Pseudotime analysis suggests that these progenitors have different differentiation trajectories, which may explain why ACT with *Nr4a3*^-/-^ effectors is more efficient. Therefore, modulation of NR4A3 activity may represent a new strategy to generate long-lived and highly functional T cells for ACT.

**One sentence summary:** NR4A3 deficiency improves anti-tumor CD8+ T cell response

## INTRODUCTION

During cancer, the persistence of the antigenic and inflammatory signals leads to progressive acquisition of the expression of inhibitory receptors (IRs) (PD-1, CD160, Lag-3, 2B4, etc.) by antigen-specific CD8^+^ T cells and loss of T cell functionality. Exhausted CD8^+^ T cells can regain functionality by blocking the interaction of IRs, such as PD-1, with their ligands. Recent studies have shown heterogeneity of CD8^+^ exhausted T cells in both chronic infection and cancer. Different states of tumor-specific CD8^+^ T cells have been identified in tumors. Among them are the less differentiated and exhausted progenitor (stem-like; TCF-1^+^ or SLAMF6^hi^Tim-3^lo^) and the terminally (TCF-1^-^ or SLAMF6^lo^Tim-3^hi^) exhausted subsets. The progenitor subset is dependent on the transcription factor TCF-1 (encoded by *Tcf7*) and will give rise to terminally Tex cells. The progenitor subset is required for successful responses to anti-PD-1/PD-L1 therapy. The importance of the progenitor subset is illustrated by the longer duration of response of melanoma patients to checkpoint blockade therapy (anti-PD-1/anti-PD-L1) when they have a larger pool of progenitor exhausted CD8^+^ T cells^1,2^.

NR4As (NR4A1, NR4A2, NR4A3) are orphan nuclear receptors that act as transcription factors to regulate the differentiation and responses of multiple immune cell types, including CD8^+^ T cells^3–6^. We have shown that NR4A3 acts as an early regulator of CD8^+^ T cell differentiation during acute response^7^. Indeed, NR4A3 deficiency in CD8^+^ T cells favors the generation of memory precursor effector cells and central memory CD8^+^ T cells while at the same time enhancing effector functions^7^. Several studies have shown that NR4As also influence CD8^+^ T cell responses during chronic responses^8–11^. It was reported that the deletion of the three members of the NR4A family (TKO) enhances the anti-tumor response of CAR-T cells used for adoptive cell therapy (ACT) of tumor-bearing mice^8^. This enhanced response is correlated with reduced CAR-T cell exhaustion and better effector functions. On the other hand, in this model, the deletion of individual NR4A had little impact on mice survival and tumor control^8^. In another study, the deletion of NR4A1 was shown to reduce CD8^+^ T cell exhaustion following chronic infection with LCMV clone 13 and following ACT of tumor-bearing mice^9^. More recently, it was shown that combined inactivation of *Nr4a3* and *Prdm1* by CRISPR in CAR-T cells increases their functionality and ability to control tumor growth in mouse pre-clinical models^11^. At the molecular level, NR4As directly contribute to the *Pdcd1* (coding for PD-1) gene transcription^8^ and restrain the accessibility of the chromatin for bZIP transcription factors^8^. Furthermore, NR4As cooperate with the transcription factor Tox, the master regulator of CD8^+^ T cell exhaustion, to induce the exhaustion program^10^.

The transcription of the three *Nr4a* genes is increased in terminally exhausted CD8 T cells and decreased following immune checkpoint blockade (ICB)^8–10,12^. Moreover, the accessibility of the chromatin containing NR4A DNA binding motif (NBRE) is lost following anti-PD-1 or anti-PD-L1 treatment^9,12^. This suggests a role for NR4A family members during the ICB response of CD8^+^ T cells. However, this was never tested experimentally.

The promotion of the memory program combined with enhanced functionality endowed *Nr4a3*^-/-^ CD8^+^ T cells^7^ with the desired attributes required for their use in ACT for the treatment of cancer. The decreased expression of *Nr4a* transcripts and DNA accessibility of NBRE motifs following ICB treatment^9,12^ further suggest that ACT with NR4A3-deficient effector CD8^+^ T cells would improve the efficacy of ACT with or without anti-PD-L1 blockade. Our results show that ACT of melanoma-bearing mice with *Nr4a3*^-/-^ effector CD8^+^ T cells provides a better tumor control than their wild-type counterpart. Moreover, the therapeutic effect observed with *Nr4a3*^-/-^ effector CD8^+^ T cells is even better than when ACT of *Nr4a3*^+/+^ effector CD8^+^ T cells is combined with anti-PD-L1 treatment. To understand how NR4A3 deficiency promotes better anti-tumor response, we have performed a scRNA-seq analysis of CD8^+^ tumor-infiltrating lymphocytes (TILs). This analysis reveals a huge heterogeneity of TIL states following ACT. The better tumor control observed with ACT of *Nr4a3*^-/-^ CD8^+^ effectors without anti-PD-L1 treatment correlates with an enrichment of TILs within the clusters that are associated with the anti-PD-L1 response of wild-type TILs. Furthermore, *Nr4a3*^-/-^ and *Nr4a3*^+/+^ effectors generate distinct progenitor populations. Monocle analysis suggests that these progenitors have different differentiation trajectories, which may explain why ACT with *Nr4a3*^-/-^ effectors is more efficient. Therefore, modulation of NR4A3 activity may represent a new strategy to generate long-lived and highly functional T cells for ACT.

## RESULTS

### ACT with NR4A3-deficient effector CD8^+^ T cells improve tumor control

Our previous report of the enhanced generation of central memory T cells, the early induction of the memory transcriptional program associated with the reduced expression of the gene signature associated with exhaustion and the increased production of cytokines by NR4A3-deficient CD8^+^ T cells^7^ led us to evaluate whether NR4A3-deficient CD8^+^ T cells will perform better in ACT to treat tumors. To test this, we decided to use *in vitro* generated effectors as our transcriptomic and ATAC-seq study have shown that NR4A3 has an early influence on the transcriptional program of CD8^+^ T cells^7^. *Nr4a3*^+/+^ and *Nr4a3*^-/-^ OT-I T cells were stimulated *in vitro* with OVA peptide-pulsed splenocytes for 48h and were then cultured with IL-2 for another 24h. These *in vitro* effectors (10^6^ cells) were adoptively transferred into mice that were previously implanted with B16-OVA tumor cells subcutaneously. A group of mice was also treated with anti-PD-L1 (Fig. 1A). ACT with NR4A3-deficient OT-I effectors was able to improve the survival of mice, while ACT with WT OT-I effectors had a smaller effect (Fig. 1B; median of survival of 32 days compared to 21). The addition of anti-PD-L1 treatment to the ACT increased the survival of mice treated with *Nr4a3^+/+^* and *Nr4a3*^-/-^ effectors (Fig. 1B; median of survival of 29 and 48 days respectively). Interestingly, ACT of *Nr4a3*^-/-^ OT-I T cells without anti-PD-L1 is even more efficient than ACT of *Nr4a3*^+/+^ OT-I T cells combined with anti-PD-L1 treatment (Fig. 1B). The therapeutic effect of PD-L1 blockade is sustained for a prolonged period after the anti-PD-L1 treatment is stopped when mice were adoptively transferred with NR4A3-deficient CD8^+^ T cells.

**Figure. 1.**
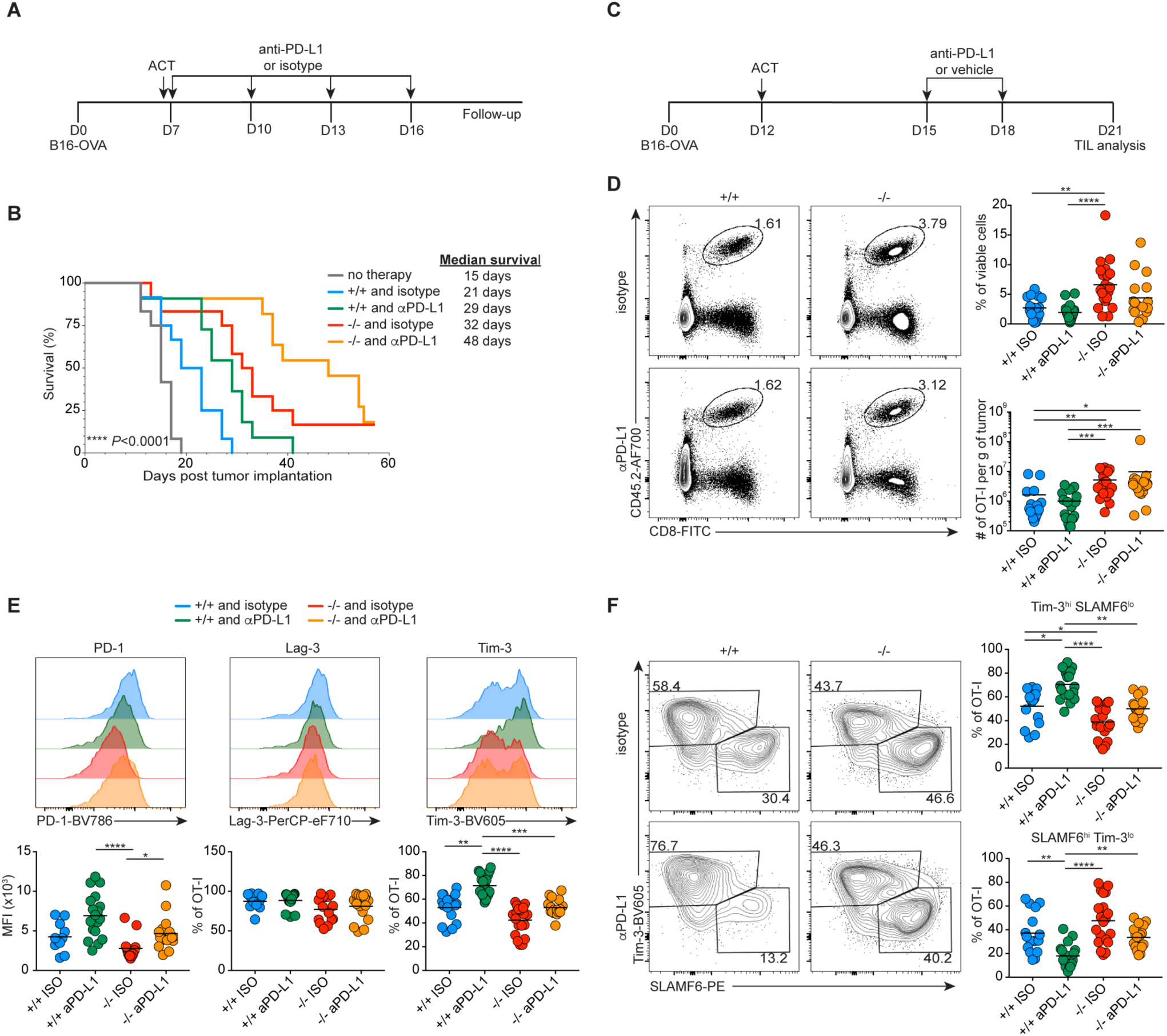
ACT with NR4A3-deficient CD8^+^ T cells improves tumor control with and without anti-PD-L1 treatment. **A.** Experimental design for combined ACT and anti-PD-L1 treatment of mice implanted with B16-OVA tumor cells. A group of mice was treated with the isotype control antibody (ISO)**. B.** Survival curve of mice implanted with B16-OVA melanoma cells under the different ACT treatments. **C.** Experimental design for TILs phenotyping after ACT combined with anti-PD-L1 treatment. At day 21 post-tumor implantation, TILs were isolated and analyzed by flow cytometry to measure OT-I cell proportion and number (**D**), inhibitory receptors expression by OT-I TILs (**E**) and progenitor (SLAMF6^hi^Tim-3^lo^) and terminal (SLAMF6^lo^Tim-3^hi^) OT-I TILs (**F**). Data are from 3 (**A, B**), or 3 to 4 (**C-F**) independent experiments. Each dot represents one mouse. Survival curves were compared using log-rank Mantel–Cox test. Kruskal-Wallis ANOVA with Dunn’s multiple comparisons was used for multiple groups comparison: \**P*<0.05, \*\**P*<0.01, \*\*\**P*<0.001, \*\*\*\**P*<0.0001.

In this setting, the combined treatment with anti-PD-L1 and *Nr4a3*^-/-^ effectors was too efficient for TIL characterization. To circumvent this, we performed ACT at day 12 post-tumor implantation, treated with anti-PD-L1 at day 15 and 18, and analyzed TILs at day 21 (Fig. 1C). This allowed to obtain measurable tumor in all experimental groups, although tumor growth was slightly reduced in mice receiving *Nr4a3*^-/-^ OT-I ACT compared to their WT counterpart (Fig. S1A and B). Furthermore, in this more aggressive model, ACT with *Nr4a3*^-/-^ effectors performed better than their WT counterparts, as seen by a higher proportion of mice in which tumors regressed between day 15 to 21 (Fig. S1C). TILs analysis of these tumors revealed increased proportion and cell numbers per gram of tumor following ACT with *Nr4a3^-/-^*OT-I effectors compared to their wild-type counterpart (Fig. 1D). However, the anti-PD-L1 treatment did not have a strong effect on the response of both *Nr4a3*^+/+^ and *Nr4a3*^-/-^ OT-I TILs (Fig. 1D). Phenotypic analysis showed that *Nr4a3*^-/-^ TILs are less exhausted than their WT counterparts as illustrated by lower level of expression of the inhibitory receptors (IRs) PD-1 and Tim-3 while no difference was observed for Lag-3 (Fig. 1E). The expression level of the PD-1 IR was increased in both *Nr4a3*^+/+^ and *Nr4a3*^-/-^ OT-I TIL following treatment with anti-PD-L1 but was still lower in NR4A3-deficient OT-I TILs. (Fig. 1E). The Tim-3^+^ population significantly increased only in *Nr4a3*^+/+^ OT-I TILs following anti-PD-L1 treatment (Fig. 1E).

ACT of OT-I *Nr4a3*^+/+^ effectors in combination with anti-PD-L1 led to an increase in the proportion of more terminally differentiated cells as shown by the increase in proportion of the SLAMF6^lo^Tim-3^hi^; PD1^+^Tim-3^+^ and CD38^+^CD101^+^ subsets (Fig. 1F and S1D-E). However, this was not observed when *Nr4a3*^-/-^ OT-I T cells were used in combination with anti-PD-L1 treatment (Fig. 1F and S1D-E). This suggests that anti-PD-L1 treatment acts by promoting the differentiation of WT progenitor/stem-like cells (SLAMF6^hi^Tim-3^lo^) into terminally differentiated effectors with a concomitant loss of progenitor/stem-like cells (Fig. 1F). However, in absence of NR4A3 expression by CD8^+^ T cells, the anti-PD-L1 treatment did not promote terminal differentiation although it enhances tumor control (Fig. 1F and S1D-E). The lack of differentiation of *Nr4a3*^-/-^ OT-I TILs into a terminally differentiated state is further illustrated by a higher proportion of SLAMF6^hi^Tim-3^lo^ OT-I TILs and by the maintenance of a PD-1^+^Tim-3^-^ phenotype with anti-PD-L1 treatment (Fig. 1F and S1E) and may explain the long-lasting beneficial effect of anti-PD-L1 blockade that is observed even after the treatment was stopped.

The transcription factors T-bet, Eomes, and TCF-1 are known to be involved in the regulation of the CD8^+^ T cell differentiation during a chronic immune response, and we previously showed that the expression of TCF-1 and Eomes are higher in NR4A3-deficient antigen-specific CD8^+^ T cells responding to an acute infection^7^. The expression of these transcription factors was similar in all experimental groups. (Fig. S1F).

Altogether, these results suggest that in absence of NR4A3, CD8^+^ T cells treated with anti-PD-L1 are less prone to terminal differentiation but are still able to efficiently control tumor growth.

### NR4A3 deficient TILs produce more TNF-α and are more polyfunctional

To determine if the increased tumor control is due to better functionality of the OT-I effectors used in the ACT, we have measured cytokine and granzyme B production following a short *ex-vivo* restimulation. The majority of OT-I TILs expressed IFN-γ with no significant differences among groups (Fig. 2A). *Nr4a3^-/-^* OT-I TILs from mice not treated with anti-PD-L1 produced more IL-2 and TNF-α compared to the *Nr4a3^+/+^* OT-I TILs from tumors treated with anti-PD-L1 (Fig. 2A). The proportion of cells co-producing IFN-γ and TNF-α was increased in *Nr4a3^-/-^* OT-I TILs from mice treated or not with anti-PD-L1 treatment when compared to *Nr4a3^+/+^* OT-I TILs with anti-PD-L1 (Fig. 2B). However, no difference was observed for IFN-γ and IL-2 co-producing cells (Fig. S2). Furthermore, anti-PD-L1 treatment did not increase the production of cytokines by both *Nr4a3*^+/+^ and *Nr4a3*^-/-^ OT-I TILs (Fig. 2A-B). Intriguingly, we also observed that *Nr4a3^+/+^* OT-I TILs under anti-PD-L1 treatment produced more granzyme B compared to the *Nr4a3^-/-^* OT-I TILs treated or not with anti-PD-L1 (Fig. 2C). Overall, *Nr4a3^-/-^* OT-I TILs were more polyfunctional, based on IFN-γ and TNF-α production, and the cytotoxicity of these cells did not increase under anti-PD-L1 treatment as it did for *Nr4a3^+/+^* OT-I TILs. These results suggest that the increased cytokine production and the increased proportions and numbers of *Nr4a3^-/-^* TILs in the tumor contribute to enhanced tumor control but may not be the only factors that explain the strong impact on mice survival observed in Fig. 1B.

**Figure. 2.**
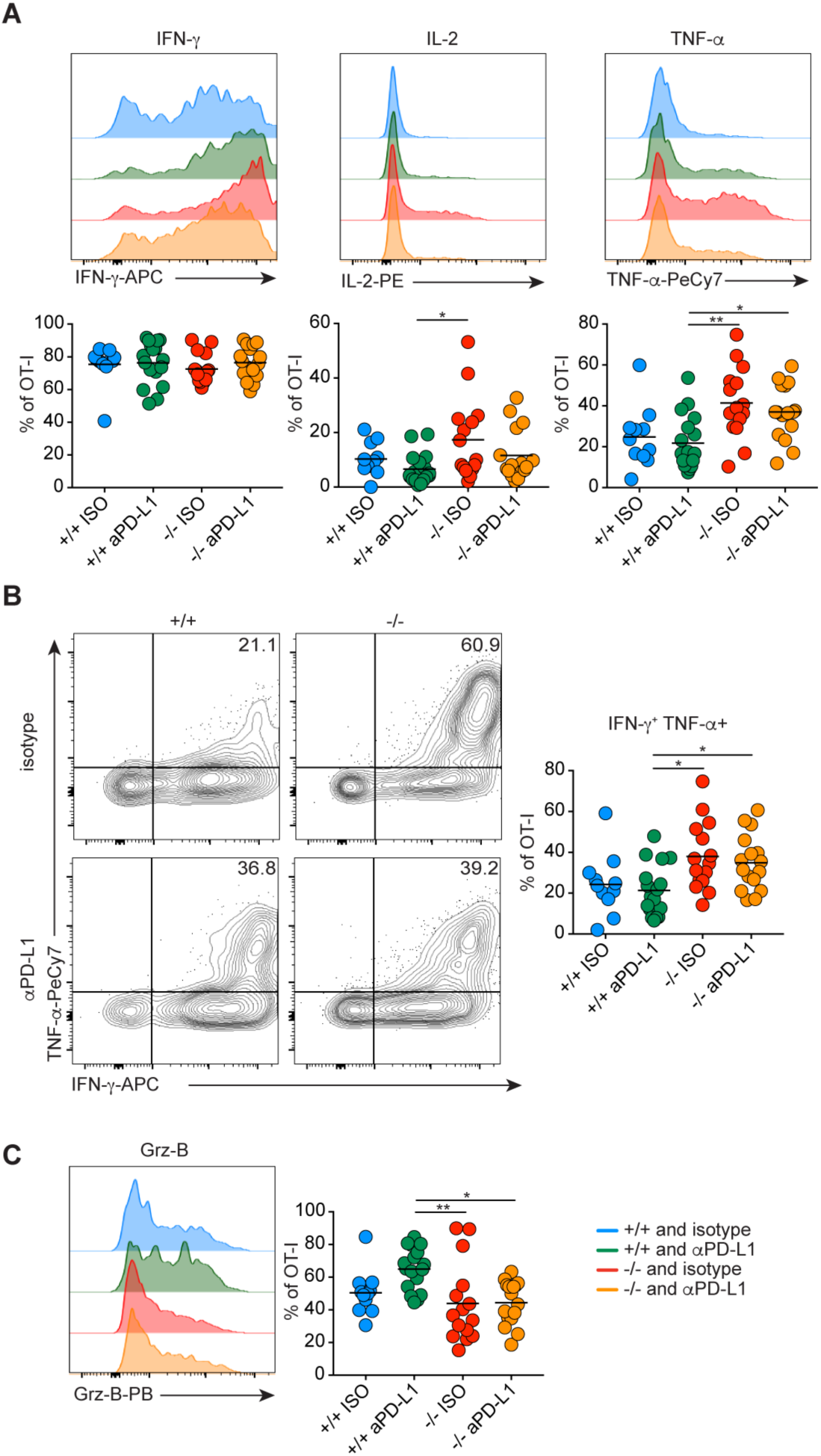
*Nr4a3*^-/-^ OT-I TILs produce more TNF-α and are more polyfunctional than *Nr4a3*^+/+^ OT-I TILs. Mice were implanted with B16-OVA melanoma cells and treated 12 days later with *Nr4a3*^+/+^ or *Nr4a3*^-/-^ *in vitro* generated OT-I effectors. Mice were treated with isotype control (ISO) or anti-PD-L1 at days 15 and 18. At day 21, tumor cell suspensions were restimulated for 5h with PMA/ionomycin for cytokine production quantification. **A-C.** Proportions of OT-I TILs producing IFN-γ^+^, IL-2^+^, or TNF-α^+^ (**A**), co-producing IFN-γ^+^ TNF-α^+^ (**B**) and granzyme-B^+^ (**C**). Data are from 3 independent experiments. Each dot represents one mouse. Kruskal-Wallis ANOVA with Dunn’s multiple comparisons was used for multiple groups comparison: \**P*<0.05, \*\**P*<0.01.

### NR4A3 deficiency confers a competitive advantage during ACT and reduces PD-1 and Lag-3 expression by WT TILs

To determine whether the reduced exhaustion and the enhanced anti-tumor efficacy of *Nr4a3^-/-^* OT-I TILs are cell-intrinsic and whether the presence of *Nr4a3^-/-^* OT-I T cells within the tumor will impact the differentiation of WT TILs, we performed competitive ACT. We treated tumor-bearing mice with a 1:1 mix of OT-I effectors (*Nr4a3^-/-^* or *Nr4a3^+/+^*) and WT OT-I/B6.SJL competitors in combination or not with anti-PD-L1 treatment (Fig. 3A). We observed a 3 to 4-fold increased accumulation of *Nr4a3^-/-^* OT-I TILs compared to their WT competitors with or without anti-PD-L1 therapy (Fig. 3B). The level of expression of the IRs, PD-1 and Lag-3 was reduced on the WT competitors when co-transferred with *Nr4a3*^-/-^ OT-I effectors but not with *Nr4a3*^+/+^ OT-I effectors (Fig. 3C). On the other hand, the *Nr4a3*^-/-^ OT-I TILs did not affect the expression of Tim-3 on the competitor WT cells (Fig. 3C). When the ACT was done with a mix of *Nr4a3*^+/+^ and *Nr4a3*^-/-^ OT-I T cells, we observed preferential differentiation of *Nr4a3*^-/-^ OT-I TILs into progenitor/stem-like cells (SLAMF6^hi^Tim-3^lo^) while the competitor *Nr4a3*^+/+^ OT-I TILs were enriched for terminally exhausted cells (SLAMF6^lo^Tim-3^hi^) (Fig. 3D). We further evaluated whether the presence of NR4A3 deficient TILs had an impact on other immune cells within the tumor microenvironment. As shown in Fig. S3, ACT with *Nr4a3*^-/-^ OT-I effectors had minimal impact on the proportion of CD4^+^, CD8^+^, Treg, B, NK, dendritic and myeloid suppressor cells. These results suggest that the effect of the deficiency of NR4A3 acts intrinsically within the OT-I TILs to improve tumor control.

**Figure. 3.**
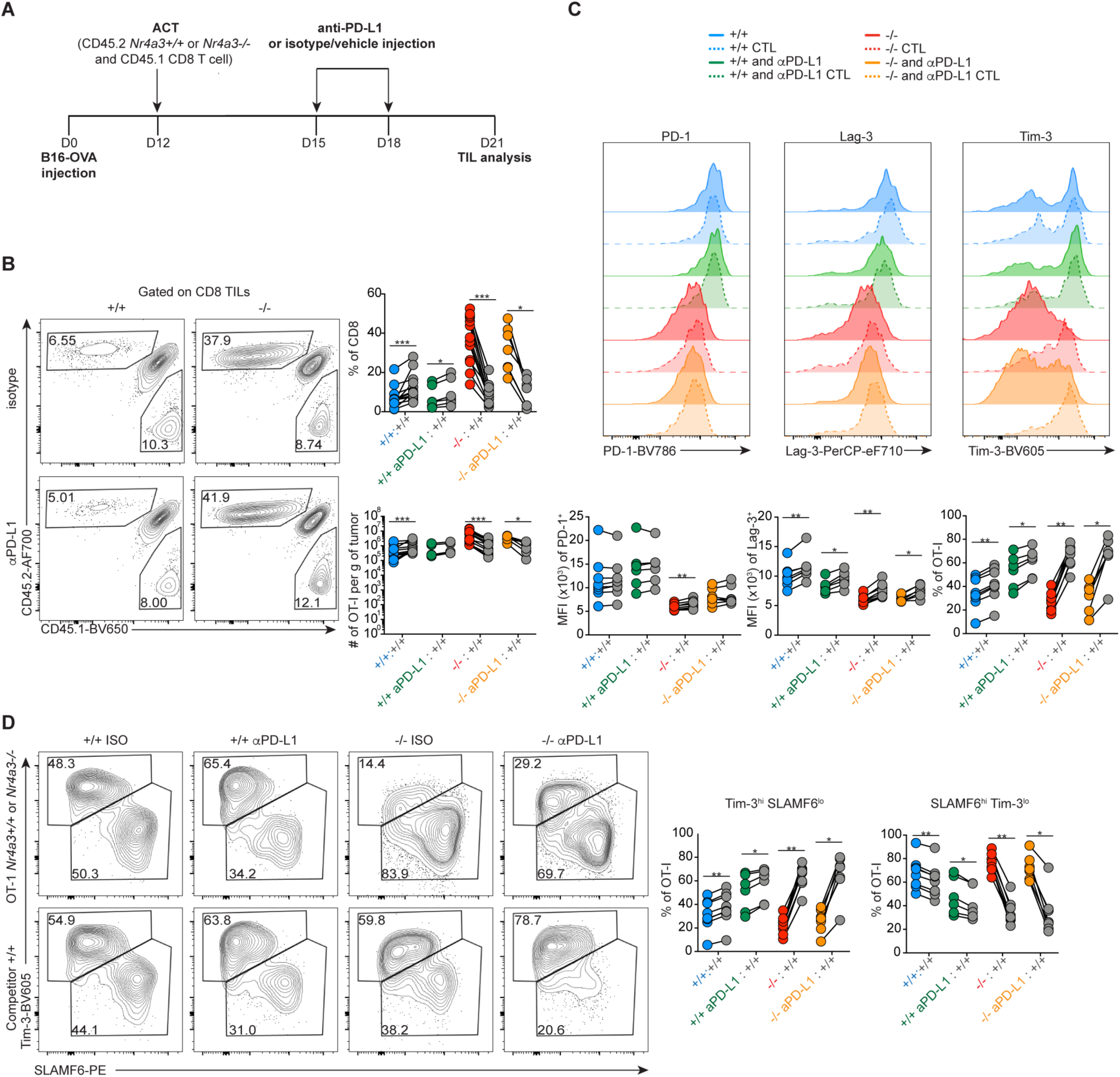
NR4A3 deficient TILs undergo less terminal exhaustion under competitive condition. **A.** Experimental design for phenotyping OT-I TILs when ACT is done in a competitive setting. At day 21 post-tumor implantation, TILs were isolated for analysis of *Nr4a3*^+/+^ or *Nr4a3*^-/-^ OT-I TILs (CD45.2^+^) and their wild-type competitors OT-I/B6.SJL TILs (CD45.1^+^). **B.** Percentage and number of OT-I TILs. **C.** Inhibitory receptor expression by OT-I TILs. **D.** Proportion of SLAMF6^lo^Tim-3^hi^ and SLAMF6^hi^Tim-3^lo^ OT-I TILs. Wild-type OT-I competitors (+/+) are shown in grey in the compilation and with dotted lines in the flow cytometry profiles. Data are from 1 experiment with 6 to 8 mice per group. Each dot represents one mouse. Paired Student’s t-test was used to compare each competitor group: \**P*<0.05, \*\**P*<0.01, \*\*\**P*<0.001.

### TIL heterogeneity following ACT with NR4A3 deficient effector CD8^+^ T cells

To define how NR4A3 deficiency enhances ACT with effector CD8^+^ T cells and to determine its impact during anti-PD-L1 treatment, we performed a single-cell RNA sequencing (scRNA-seq) analysis on sorted *Nr4a3^+/+^* and *Nr4a3^-/-^*OT-I TILs with or without anti-PD-L1 therapy on day 21 (as done for TIL characterization (Fig. 1C-F)). Sequenced TILs transcriptomes were grouped in 15 clusters (C0 to C14) and we observed a different cell distribution of *Nr4a3^+/+^*and *Nr4a3^-/-^* TILs with or without anti-PD-L1 treatment on their UMAP projection (Fig. 4A).

**Figure. 4.**
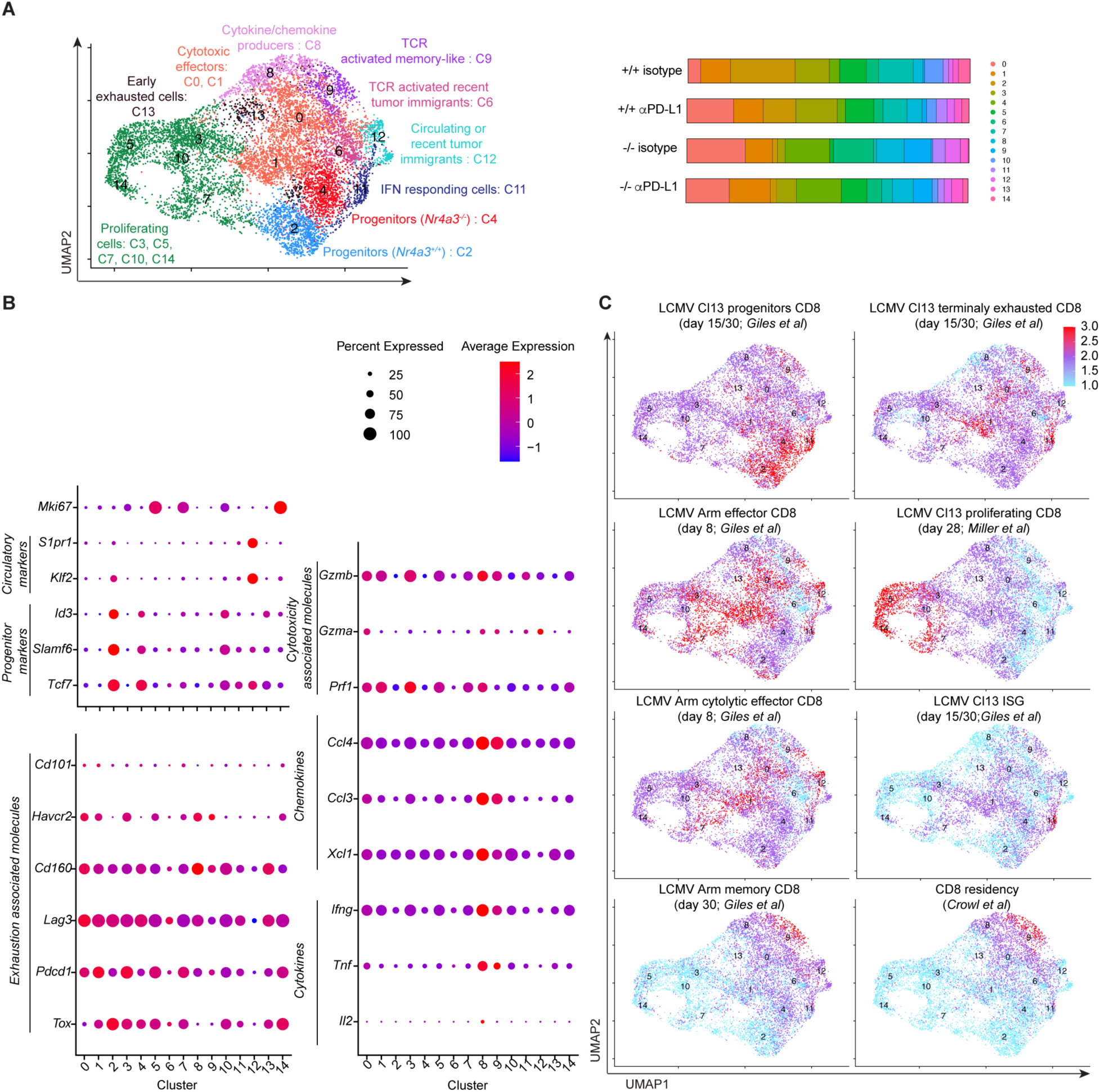
TIL heterogeneity following ACT with effector CD8^+^ T cells. Mice were implanted with B16-OVA melanoma cells and treated 12 days later with *Nr4a3*^+/+^ or *Nr4a3*^-/-^ *in vitro* generated OT-I effectors. Anti-PD-L1or isotype control antibody treatment was done on days 15 and 18. At day 21, OT-I TILs were isolated to perform scRNA-seq analysis. **A.** UMAP from scRNA-seq with annotation of functional clusters (left) and *Nr4a3*^+/+^ or *Nr4a3*^-/-^ TILs distribution among the clusters (right). **B**. Level of expression of selected genes by cluster. **C**. Gene signature projections of different CD8^+^ T cell subsets on the UMAP. Data are from one scRNA-seq experiment where 3 independent biological samples were pooled for each treatment condition.

The identity of the 15 clusters was defined based on gene expression (markers), gene signatures from published studies^2,13–16^ as well as gene ontology (GO) and GSEA gene enrichments (Fig. 4A). We defined clusters 2 and 4 as progenitors as these cells are enriched for transcription of *Tcf7*, *Slamf6,* and *Id3* combined with a low level of *Havcr2* (Fig. 4B). These two clusters are enriched for multiple gene signatures of progenitor/stem-like CD8^+^ T cells isolated from tumors (Fig. S4A) or chronic LCMV infection (Fig. 4C). As expected for progenitors, clusters 2 and 4 do not show enrichment for terminally exhausted gene signatures (Fig. 4C *Giles et al.*^13^ and Fig. S4A *Siddiqui et al.*^2^). These two progenitor clusters are almost exclusively distributed within *Nr4a3*^+/+^ and *Nr4a3*^-/-^ OT-I TILs, with cluster 2 being enriched in the former and cluster 4 in the latter (Fig. 4A and Table I).

**Table I.** scRNA-seq TILs distribution (%) among the 15 clusters.

| Cluster | *Nr4a3*<sup>+/+</sup> ISO | *Nr4a3*<sup>+/+</sup> αPD-L1 | *Nr4a3*<sup>-/-</sup> ISO | *Nr4a3*<sup>-/-</sup> αPD-L1 |
| --- | --- | --- | --- | --- |
| 0 | 4,5 | 16,5 | 20,7 | 15,4 |
| 1 | 10,6 | 10,5 | 9,6 | 14,5 |
| 2 | 22,9 | 11,5 | 1,8 | 2,3 |
| 3 | 12,2 | 14,7 | 2,7 | 7,0 |
| 4 | 3,6 | 2,9 | 15,8 | 15,9 |
| 5 | 9,5 | 10,1 | 1,3 | 9,1 |
| 6 | 4,4 | 3,2 | 14,2 | 5,2 |
| 7 | 12,3 | 8,5 | 1,1 | 3,7 |
| 8 | 3,4 | 5,4 | 9,8 | 7,4 |
| 9 | 0,6 | 1,5 | 9,6 | 6,7 |
| 10 | 6,6 | 3,5 | 0,5 | 2,1 |
| 11 | 1,9 | 3,8 | 4,4 | 2,2 |
| 12 | 1,6 | 2,6 | 5,3 | 2,6 |
| 13 | 2,0 | 2,6 | 2,8 | 3,9 |
| 14 | 4,1 | 2,6 | 0,3 | 1,9 |

Cluster 12 contains TILs with the characteristic of circulating cells and recent tumor immigrants. Although the TILs within cluster 12 are enriched for the expression of the progenitor-associated genes *Tcf7* and *Slamf6,* their expression of *Klf2 and S1pr1* suggests that they represent TILs that are circulating and have recently entered into the tumors (Fig. 4B). This is reinforced by their low level of transcription of exhaustion-associated genes (*Tox*, *Pdcd1*, *Lag3*, *Cd160, Havcr2*, *Cd101*) (Fig. 4B). Cluster 12 cells are also enriched for the gene signature of naïve versus exhausted CD8^+^ T cells (GSE9650; *Padj* = 5.85*10^-6^ and NES = 2.85) and for the gene signature of central memory compared to effector memory CD8^+^ T cells (Fig. S4A). Altogether, this indicates that cluster 12 contains OT-I effector cells that have recently entered in the tumors and as such are less exhausted.

Clusters 6 and 9 are both enriched for TCR signaling gene signature. Cluster 6 is enriched for the GO pathway associated with TCR binding (GO: 0042608; FDR = 5.00*10^-3^) and KEGG TCR signaling pathway (KEGG:04660; FDR = 2.83*10^-6^). Similarly, cluster 9 is enriched for the GO pathway associated with the TCR signaling pathway (GO: 0050852; FDR = 1.82*10^-8^), KEGG TCR signaling pathway (KEGG:04660; FDR = 2.11*10^-7^) and regulation of the TCR signaling pathway (GO: 0050856; FDR = 1.33*10^-2^). TILs within both clusters are not enriched for exhaustion-associated genes (Fig. 4B). Although both clusters contain non-exhausted TCR-activated cells, only cluster 9 showed enrichment for effector, memory and residency gene signatures (Fig. 4C). Cluster 6 probably represents TCR-activated recent tumor immigrants as they do not transcribe genes associated with effector functions (*Tbx21*, *Zeb2, Batf*, *Prdm1*), early activation markers (*Cd69*, *Il25ra*) and TCR induced genes (*Nr4a1, Nr4a2*, *Irf4*) (Fig. S4B). On the other hand, cluster 9 can be defined as TCR-activated memory-like cells as these TILs are enriched in memory and residency gene signatures (Fig. 4C) and actively transcribes early activation (*Cd69*, *Il25ra*) and TCR-induced (*Nr4a1, Nr4a2*, *Irf4*) genes (Fig. S4B). Interestingly, these two clusters are overrepresented in *Nr4a3^-^*^/-^ TILs (Fig. 4A and Table I).

Clusters 0 and 1 are defined as cytotoxic effectors. These TILs are enriched for the gene signature of effectors and cytotoxic cells (Fig. 4C; *Giles et al.*^13^). The clusters 0 and 1 are also enriched for gene signature from effectors compared to exhausted CD8^+^ T cells (GSE9650; Cluster 0: *Padj* = 1.66*10^-8^ and NES = 3.58; Cluster 1: *Padj* = 1.01*10^-12^ and NES = 4.19). These two clusters also transcribed genes encoding effector molecules (*Gzmb*, *Gzma*, *Prf1* and *Ifng*; Fig. 4B)) and transcription factors associated with effector differentiation (*Tbx21* and *Zeb2*; Fig. S4B). The TILs within these clusters also express some exhaustion-associated genes (*Tox*, *Pdcd1*, *Lag3*, *Cd160, Havcr2*) (Fig. 4B) and show enrichment for exhausted CD8^+^ T cells gene signature (Fig. 4B *Giles et al.*^13^ and Fig. S4A *Siddiqui et al.*^2^) which suggests that these cells may start to be exhausted but have still potent effector functions.

Cluster 8 is also an effector cluster highly enriched for cytokine and chemokine production. This cluster contains TILs transcribing high levels of genes encoding for cytokines (*Il2*, *Tnf, Ifng*), chemokines (*Xcl1*, *Ccl3*, *Ccl4*), and cytotoxicity-associated molecules (*Prf1*, *Gzma*, *Gzmb*) (Fig. 4B). These cells are also enriched in GO pathways associated with chemokine activity (GO:0008009; FDR = 3.09*10^-3^), regulation of cytokine production (GO:0001817; FDR = 7.07*10^-14^), positive regulation of cytokine production (GO:0001819; FDR = 6.19*10^-12^), cytokine production involved in immune response (GO:0002367; FDR = 7.50*10^-4^) as well as the negative regulation of cytokine production (GO:0001818; FDR = 1.89*10^-3^). Although this cluster contains cells that produced high levels of effector molecules, it shows less enrichment for the effector and cytotoxic gene signature than clusters 0 and 1 (Fig. 4B). Cluster 8 is also enriched with TILs expressing transcription factors associated with effector functions such as *Tbx21* and *Zeb2* (Fig. S4B) while not expressing the exhaustion transcription factor *Tox* (Fig. 4B).

Clusters 3, 5, 7, 10, and 14 are defined as proliferating cells as they show enrichment in cell proliferation-associated GO pathways. The most significantly enriched GO pathways within these clusters are: mitotic cell cycle (GO:0000278 and GO:1903047), cell cycle (GO:0007049), cell cycle process (GO:0022402), cell division (GO:0051301) and DNA replication (GO:0006260). The TILs in these clusters show a high level of transcription of *Mki67* (Fig. 4B), as well as enrichment of the gene signature of proliferating cells (Fig. 4C) and cell cycle (Fig. S4A). These different clusters of proliferating cells can be distinguished into proliferating progenitors (cluster 10) and effectors (clusters 3, 5, 7 and 14) (Fig. 4B-C). The heterogeneity among the proliferating cell populations illustrates the complexity of TIL populations during an anti-tumor immune response.

Cluster 11 contained TILs responding to IFN as illustrated by their enrichment for the expression of *Stat1, Stat2, Irf1, Irf2, Irf7,* and *Bst2* (Fig. S4B) and the interferon-stimulated genes (ISG) signature (Fig. 4C). Cluster 13 shows enrichment for the effector and cytotoxic CD8^+^ T gene signatures but to a lesser extent than other clusters (Fig. 4C). This observation combined with the expression of *Lag3* and *Cd160* but not of *Havcr2* (Fig. 4B) suggests that the TILs within cluster 13 are at an early stage of T cell exhaustion.

### NR4A3 deficiency induces a transcriptional program of anti-PD-L1 responding TILs

The distribution of TILs within the 15 clusters is very different depending on their genotypes (Fig. 5A and Table I). The *Nr4a3*^-/-^ TILs are more abundant on the right side of the scRNA-seq UMAP (C0, C1, C4, C6, C8, C9, C11, C12) when compared to the *Nr4a3^+/+^*TILs that are more distributed on the left side (C1, C2, C3, C5, C7, C10, C14) (Fig. 5A). The anti-PD-L1 treatment shifted the *Nr4a3^+/+^* TILs distribution to the right side of the UMAP where *Nr4a3*^-/-^ TILs without anti-PD-L1 treatment are already more prevalent (Fig. 5A). This suggests that ACT with NR4A3-deficient effectors allows for better tumor control due to a transcriptomic signature similar to the one induced by anti-PD-L1 treatment. Another striking difference within *Nr4a3*^+/+^ and *Nr4a3*^-/-^ TILs, with and without anti-PD-L1 treatment, is the observation that the *Nr4a3*^+/+^ and *Nr4a3*^-/-^ TILs with a progenitor signature are found in different cluster according to their genotype (Fig. 5A and Table I).

**Figure 5.**
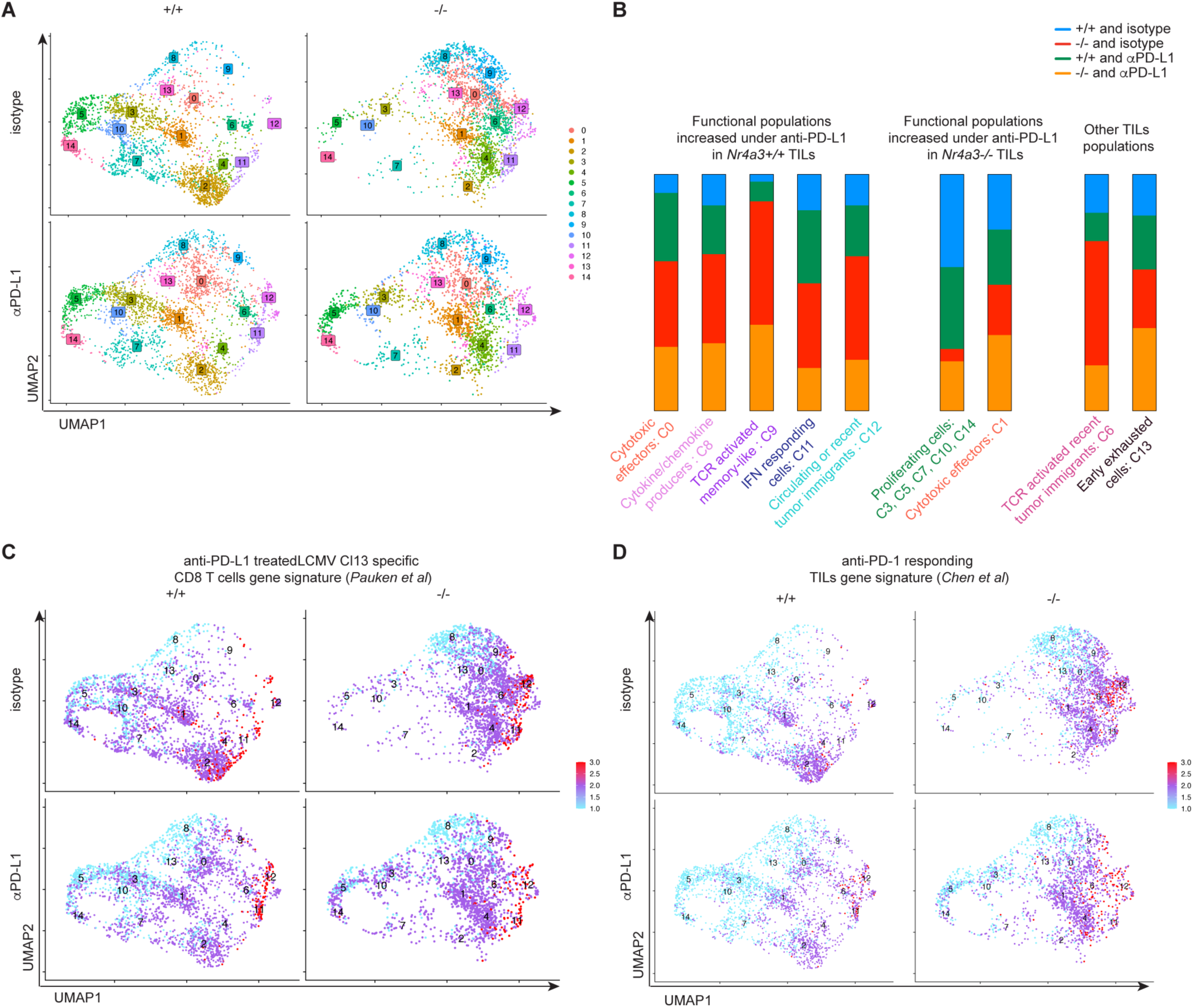
NR4A3 deficiency induces a transcriptional program of anti-PD-L1 responding TILs. A. UMAP distribution of the 15 scRNA-seq clusters in function of TILs genotype and treatment conditions. **B.** TILs distribution in functional clusters by treatment condition. **C**. Gene signature projection of anti-PD-L1 treated CD8^+^ T cells on the scRNA-seq UMAP of *Nr4a3*^+/+^ or *Nr4a3*^-/-^ TILs under the anti-PD-L1 or the isotype control treatment. **D**. Immune checkpoint blockade (ICB) responder gene signature projection on the UMAP of *Nr4a3*^+/+^ or *Nr4a3*^-/-^ TILs under the anti-PD-L1 or the isotype control treatment.

To explain the increased tumor control induced by ACT with *Nr4a3^-/-^*effectors CD8^+^ T cells, we studied TIL population dynamics associated with the different ACT regimens. Following ACT with *Nr4a3*^+/+^ effectors, the anti-PD-L1 treatment induces an increase in effectors (3.7x C0), cytokine and chemokine producers (1.6x C8), TCR-activated memory-like effectors (2.6x C9), IFN responding TILs (2.0x C11) and circulating or recent tumor immigrants (1.6x C12) (Fig. 5B and Table I). These same clusters are also more abundant following ACT with *Nr4a3*^-/-^ effectors in the absence of anti-PD-L1 treatment (Fig. 5A-B). Indeed, when we compare the *Nr4a3^-/-^*TILs to *Nr4a3^+/+^* TILs treated with the isotype control antibody, we observe increased proportions of effectors (4.6x C0), cytokine and chemokine producers (2.9x C8), TCR activated memory-like effectors (16.3x C9), IFN responding TILs (2.4x C11) and circulating or recent tumor immigrants (3.4x C12) (Fig. 5B and Table I). Overall, the populations that are induced by the anti-PD-L1 treatment in the wild-type setting are already enriched in NR4A3 deficient TILs without PD-1/PD-L1 blockade, and these clusters are the ones containing TILs with effector functions. This helps to explain why ACT with *Nr4a3*^-/-^ effectors is as efficient as ACT with *Nr4a3*^+/+^ effectors combined with PD-L1 blockade to control tumor growth. The *Nr4a3^-/-^* TILs under the anti-PD-L1 treatment undergo different cluster dynamics, where the anti-PD-L1 treatment mainly induces an increase in cycling cells (2.6x C3; 7.1x C5; 3.4x C7; 3.8x C10; 5.8x C14) and some effectors (Fig. 5B and Table I). The increase in the proliferation rate of TILs may explain how anti-PD-L1 treatment enhanced the efficacy of ACT with *Nr4a3*^-/-^ effector CD8^+^ T cells.

To study the impact of NR4A3 and anti-PD-L1 treatment on the TIL transcriptional changes, we evaluated the transcriptional differences among the different treatment conditions between cells of the same cluster. The highest number of differentially expressed genes was observed when comparing *Nr4a3^+/+^*with *Nr4a3^-/-^* TILs under isotype control treatment, with a similar trend, although at a lower level, for these TILs comparison under the PD-L1 blockade (Fig. S5). The lowest number of differentially expressed genes within each cluster was observed when comparing *Nr4a3^-/-^* TILs treated or not with anti-PD-L1 (Fig. S5). *Nr4a3^+/+^* TILs also undergo greater changes under anti-PD-L1 treatment than *Nr4a3^-/-^* TILs. Moreover, the number of differentially expressed genes decreased between *Nr4a3^+/+^* and *Nr4a3^-/-^* TILs under the anti-PD-L1 treatment, which suggest that PD-L1 blockade reduces transcriptional differences among *Nr4a3^+/+^* and *Nr4a3^-/-^* TILs (Fig. S5).

To illustrate the changes that occur under the anti-PD-L1 treatment, we projected on our UMAP an available gene signature of TILs following anti-PD-L1 treatment^12^ (Fig. 5C). This gene signature is highly enriched on the right side of the UMAP, where we mainly found *Nr4a3^-/-^* TILs (Fig. 5C). The most enriched populations for this gene signature are IFN responding TILs (C11) and circulating or recent tumor immigrants (C12) which proportions are increased under anti-PD-L1 treatment in *Nr4a3^+/+^*TILs and in *Nr4a3^-/-^* TILs without treatment (Fig. 5C and Table I).

To determine if the transcriptional landscape of *Nr4a3^-/-^* TILs could explain why they are better at controlling tumor growth, we projected an immune checkpoint blockade (ICB) responder gene signature^17^ on our data (Fig. 5D and S5B). Overall, we observed an enriched responder gene signature on the right side of the UMAP in the progenitor clusters (C2 and C4) and in clusters where *Nr4a3^-/-^* TILs are enriched (Fig. 5D). This responder gene signature is more abundant in *Nr4a3^-/-^* TILs with or without anti-PD-L1 treatment (Fig. 5D) when compared to the *Nr4a3^+/+^* conditions.

In conclusion, *Nr4a3^+/+^* and *Nr4a3^-/-^* TILs functional populations with or without the anti-PD-L1 treatment are distinct in their proportions and transcriptional profile inside each cluster. *Nr4a3^-/-^*TILs are transcriptionally predicted to be better responders during an anti-tumor response and already have an anti-PD-L1-treated like transcriptional profile without PD-L1 blockade.

### NR4A3 expression influences the differentiation and transcriptional profile of TILs

Our scRNA-seq identified two different clusters for *Nr4a3^+/+^*and *Nr4a3^-/-^* progenitors. These clusters were enriched in gene signatures of progenitors (Fig. 4C and S4A) and in *Tcf7* expression (Fig. S6). Cluster 2 contains mainly *Nr4a3^+/+^*TILs, while cluster 4 is associated with *Nr4a3^-/-^* TILs (Fig. 5A and Table I). The different clustering of *Nr4a3^+/+^*and *Nr4a3^-/-^* progenitors suggests different biological proprieties. To illustrate this, we performed a pseudotime analysis to predict the differentiation trajectory of the two progenitor populations during the anti-tumor response with and without anti-PD-L1 treatment (Fig. 6A). To initiate the predicted pathway of differentiation in the pseudotime analysis, we used as starting point the cluster 2 for the *Nr4a3^+/+^* TILs and cluster 4 for the *Nr4a3^-/-^* TILs. This analysis allowed us to observe that the predicted pathways of differentiation of *Nr4a3^+/+^*and *Nr4a3^-/-^* progenitors are distinct (Fig. 6A). The *Nr4a3^+/+^* progenitors differentiate into cycling cells (C3, C5, C7, C10, C14) before giving rise to cells with effector functions. Under anti-PD-L1 treatment, the differentiation pathway of the *Nr4a3^+/+^*progenitors (C2) is changed, they first become effectors (C1) before differentiating into cycling cells (C3, C5, C7, C10, C14) that differentiate further into cells with effector functions. Furthermore, under anti-PD-L1 treatment *Nr4a3*^+/+^ TILs further differentiate into TCR-activated memory-like effectors (C9) and IFN-responding cells (C11). The *Nr4a3^-/-^* progenitors have a different pathway of differentiation, from progenitors (C4), they directly differentiate into effectors (C0, C1) and into TCR-activated recent tumor immigrants (C6), TCR-activated memory-like effectors (C9) and cytokine/chemokine producers (C8). Under anti-PD-L1 treatment, the *Nr4a3^-/-^* progenitors differentiation change, progenitors (C4) give rise to cycling cells (C3, C5, C7, C10, C14) that are very rare without PD-L1 blockade. Another striking difference is that under anti-PD-L1 treatment, the proportion of *Nr4a3*^-/-^ progenitors does not change while their wild-type counterpart decreased (Fig. 6B and Table I). This is reminiscent of what we have observed by flow cytometry using SLAMF6/Tim-3 stainings (Fig. 1F). Altogether, this suggests that the different proprieties of *Nr4a3^+/+^* and *Nr4a3^-/-^* progenitors impact their differentiation path and suggests that *Nr4a3^-/-^* progenitors are able to directly differentiate into effectors and cells populations that are enriched under the anti-PD-L1 treatment more rapidly than *Nr4a3^+/+^* progenitors, which need to proliferate before differentiating into effectors.

**Figure 6.**
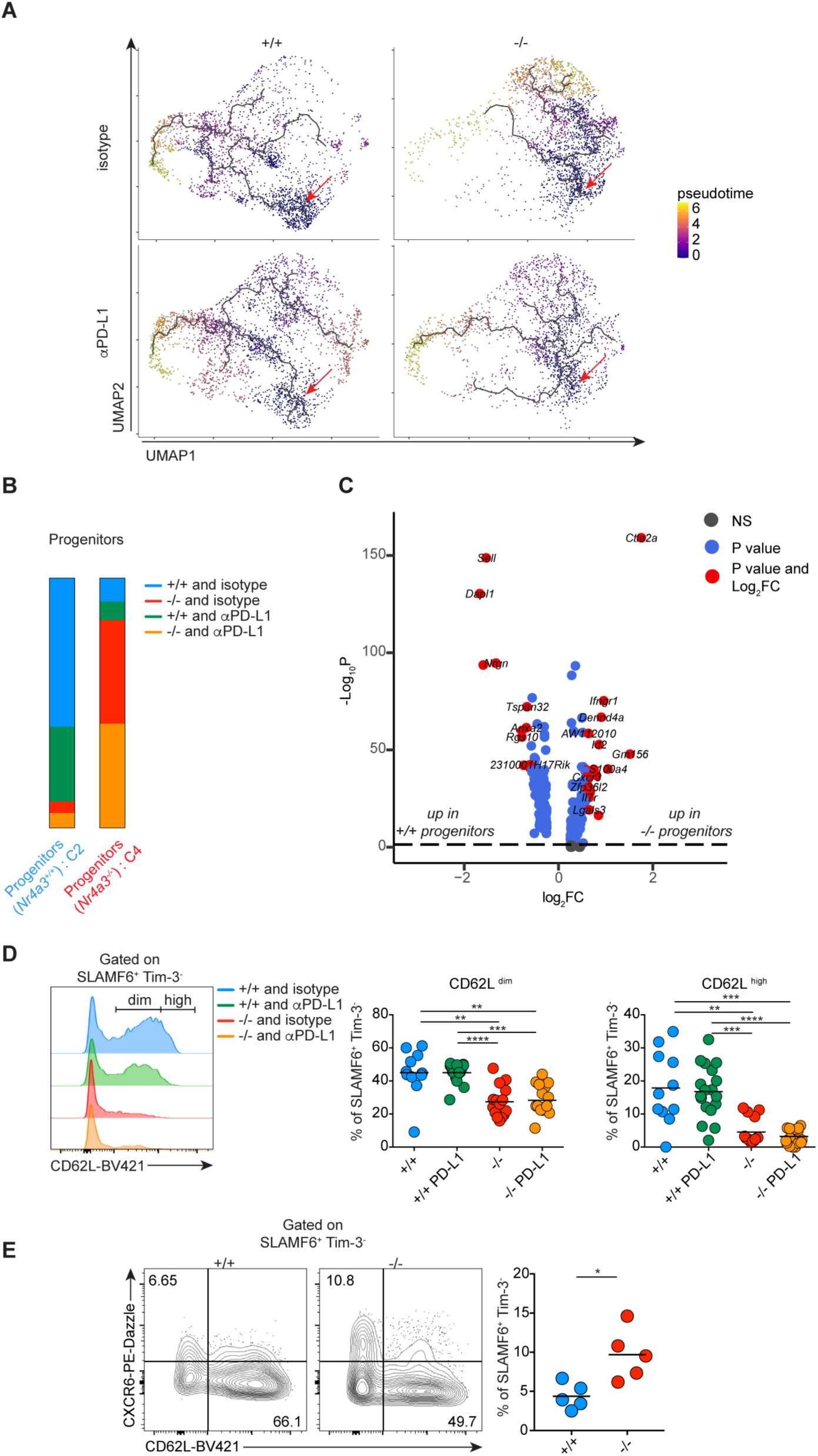
NR4A3-deficiency in CD8^+^ T cells induces the differentiation of distinct progenitor population. **A.** Pseudotime analysis of the differentiation trajectory of *Nr4a3*^+/+^ or *Nr4a3*^-/-^ TILs under anti-PD-L1 or isotype control treatment. **B.** Progenitor cell distributions in clusters 2 and 4 by treatment condition. **C.** Differentially expressed genes among *Nr4a3*^-/-^ (cluster 4) or *Nr4a3*^+/+^ (cluster 2) progenitors. **D-E.** CD62L (**D**) and CXCR6 (**E**) expression by OT-I by *Nr4a3*^+/+^ or *Nr4a3*^-/-^ TIL progenitors (SLAMF6^hi^Tim-3^lo^). Data are from 3 (**D**) or one (**E**) phenotyping TILs experiment (day 21) where each point represents one mouse.

To better characterize the *Nr4a3^+/+^* and *Nr4a3^-/-^* progenitors, we compared their gene expression profiles. We identified 273 differentially expressed genes (*Padj*. <0.05) between the *Nr4a3*^+/+^ and *Nr4a3*^-/-^ TIL progenitors. Among them, 22 show more than 1.5-fold enrichment in *Nr4a3^-/-^* progenitors and 12 in *Nr4a3^+/+^*progenitors (Fig. 6C). Among these genes, we searched for extracellular molecules coding genes to be able to find markers that will allow us to distinguish *Nr4a3^+/+^* and *Nr4a3^-/-^* progenitors. *Ifngr1, Cxcr6, Emb, Il7r, and Jaml* gene expression is higher in *Nr4a3^-/-^* progenitors, while *Dapl1, Sell,*

*Nrgn, Itga4* expression is increased in *Nr4a3^+/+^* progenitors. We then evaluated whether these markers allow us to distinguish by flow cytometry the wild-type progenitors from the NR4A3-deficient ones. The only two markers for which we observed different levels of protein expression between *Nr4a3*^+/+^ and *Nr4a3*^-/-^ progenitors are CD62L (encoded by *Sell*) and CXCR6 (Fig. 6D-E and data not shown). CD62L expression is higher on *Nr4a3^+/+^* progenitors (Fig. 6D) and CXCR6 on *Nr4a3^-/-^* progenitors (Fig. 6E).

We defined *Nr4a3^+/+^* and *Nr4a3^-/-^* progenitors as cells with a distinct transcriptome that could differentiate by different predicted pathways in order to sustain the anti-tumor immune response. Furthermore, *Nr4a3^+/+^* and *Nr4a3^-/-^* progenitors are transcriptionally distinct, the *Nr4a3^+/+^* expressing higher *Sell* gene and CD62L protein, whereas *Nr4a3^-/-^* progenitors express higher *Cxcr6* gene and CXCR6 protein.

## DISCUSSION

Our results show that NR4A3-deficient CD8^+^ T cells have enhanced ability to control tumor growth in an ACT mouse model of melanoma and that this therapeutic effect is further enhanced by PD-L1 blockade. This better tumor control following ACT with *Nr4a3*^-/-^ effector CD8^+^ T cells is associated with an increased proportion of *Nr4a3*^-/-^ CD8^+^ TILs within the tumors and a decrease in their exhaustion and terminal differentiation. scRNA-seq analysis of TILs further reveals: i) the presence of distinct subsets of progenitors following ACT with *Nr4a3*^+/+^ and *Nr4a3*^-/-^ effector CD8^+^ T cells; ii) the enrichment of *Nr4a3*^-/-^ TILs within the clusters that are associated with the anti-PDL-1 response of wild-type TILs. Altogether, this suggests that following ACT, *Nr4a3*^-/-^ effectors differentiate into TILs with a transcriptional profile of ICB-treated cells even without PD-L1 blockade.

One striking observation that we have made, is that ACT with *Nr4a3*^-/-^ effector CD8^+^ T cells is better than ACT with *N4ra3*^+/+^ effectors combined with anti-PD-L1 therapy. This is probably explained by the differentiation of *Nr4a3*^-/-^ effector CD8^+^ T cells into TIL subsets having the transcriptional characteristics of wild-type TILs under PD-L1 blockade. Indeed, *Nr4a3*^-/-^ TILs are enriched in clusters associated with effector functions and these clusters are the ones that increased following anti-PD-L1 treatment of wild-type cells. At the molecular level, the reduced exhaustion and enhanced effector functions of *Nr4a3*^-/-^ TILs could be a direct consequence of the reported ability of NR4As to reduce the accessibility of chromatin containing binding motifs for bZIP transcription factors^9^, as bZIP transcription factors are known to control T cell activation and effector functions^18–22^. Furthermore, c-Jun, a bZIP transcription factor overexpression was shown to reduce T cell exhaustion and to enhance effector functions of CAR T cells^23^. Further studies are required to define whether the better anti-tumor activity of *Nr4a3*^-/-^ effector CD8^+^ T cells is the result of enhanced binding of bZIP transcription factors to DNA. The better tumor control following ACT with *Nr4a3*^-/-^ effectors compared to *Nr4a3*^+/+^ effectors combined with anti-PD-L1 treatment is in line with previous observations that have shown that anti-PD-L1 treatment decreases transcription of the *Nr4a* genes and the chromatin accessibility of NR4As. Therefore, we would like to suggest that one of the main mechanisms of action of anti-PD-L1 treatment is via the regulation of NR4A activity. However, anti-PD-L1 treatment does not solely acts through the modulation of NR4A3 activity as PD-L1 blockade enhances the therapeutic efficacy of ACT with *Nr4a3*^-/-^ effector CD8^+^ T cells.

Although anti-PD-L1 promotes better tumor control following ACT with *Nr4a3*^-/-^ effectors, the mechanism by which it does so seems to be very different than ACT with wild-type cells. Indeed, PD-L1 blockade induces the differentiation of *Nr4a3*^+/+^ TIL progenitors (SLAMF6^hi^ Tim-3^lo^) into more exhausted TILs (SLAMF6^lo^Tim-3^hi^), while this is not observed with *Nr4a3*^-/-^ TILs. Furthermore, most TIL subsets identified with the scRNA-seq do not change in proportion in absence of NR4A3, while they change significantly in the wild-type setting. Furthermore, anti-PD-L1 treatment induces significant changes in the transcriptome of the different *Nr4a3*^+/+^ TIL clusters, while changes are minimal in absence of NR4A3. In agreement with our flow cytometry data, the proportion of cells in the cluster of *Nr4a3*^-/-^ progenitors does not change with PD-L1 blockade, while it decreases for *Nr4a3*^+/+^ TILs. This raised the question of how NR4A3-deficient TILs control better the tumor growth under anti-PD-L1 treatment. It is likely that maintenance of stemness in *Nr4a3*^-/-^ progenitor TILs promotes better tumor control by allowing constant fueling of effectors while maintaining themselves. On the other hand, it is possible that following anti-PD-L1 treatment, *Nr4a3*^-/-^ progenitors are constantly recruited from secondary lymphoid organs. This fits with our previous observation that NR4A3-deficient CD8^+^ T cells generated more central memory CD8^+^ T cells during acute infection^7^. The different behavior of *Nr4a3*^+/+^ and *Nr4a3*^-/-^ progenitors following anti-PD-L1 treatment probably reflects the fact that these progenitors have a distinct transcriptional profile. Indeed, *Nr4a3*^+/+^ and *Nr4a3*^-/-^ progenitors are enriched within different clusters. We have identified some cell surface molecules that distinguish the *Nr4a3*^+/+^ and *Nr4a3*^-/-^ progenitors; CD62L is expressed at higher level on wild-type progenitors, while CXCR6 is higher on NR4A3-deficient progenitors. The differential expression of these molecules could have functional importance as CD62L is known to define stem-like TILs^24^ while CXCR6^+^ TILs are recruited by IL-15 presented by CCR7^+^ DCs to promote long-term survival^25^. Further studies should reveal the functional difference and differentiation pathway associated with these distinct progenitor subsets.

Our results on ACT of cancer also highlight that deficiency in only one member of the NR4A family is sufficient to provide better therapeutic efficacy of CD8^+^ effectors. Interestingly, another group recently reported that ACT with *Nr4a1*^-/-^ CD8^+^ T cells led to better tumor control than their WT counterpart^9^. This suggests that the different NR4A family members can have dominant effect in the context of anti-tumor response. Further studies should reveal whether they do so via a similar mechanism. In contrast, another group recently reported that the deletion of the three NR4A family members was required to enhance the action of chimeric-antigen receptor (CAR) expressing CD8^+^ T cells^8^. Furthermore, a recent study showed that NR4A3 deletion in human CAR-T cells was not sufficient to enhance tumor control but that the co-deletion of *Nr4a3* and *Prdm1* (coding for Blimp-1) is required to decrease exhaustion and improve tumor control^11^. One possible explanation for these differences could relate to the stronger antigenic signal that may lead to more severe exhaustion of CAR-expressing TILs, which might not be overcome by the deletion of only one NR4A member in CD8^+^ TILs.

Altogether, our results suggest that the modulation of NR4A3 activity represents a promising avenue for the generation of better T-cell products for ACT of cancer patients and improving the efficacy of anti-PD-1/PD-L1 treatment of cancer patients.

## MATERIALS AND METHODS

### Mice

*Nr4a3^-/-^* and *Nr4a3^+/+^*, OT-I (*Rag1^-/-^*)^26^, B6.SJL, C57BL/6 and CD45.1.2 (F1 of B6.SJL x C57BL/6) mice were all bred at the Maisonneuve-Rosemont Hospital Research Center facility. *Nr4a3^-/-^* mice were kindly provided by Dr. Orla M. Conneely^27^ and were backcrossed for at least 10 generations to C57BL/6 mice. These mice were then crossed to the OT-I mouse strain to obtain OT-I *Nr4a3^+/+^* (*Rag1^-/-^*) and OT-I *Nr4a3^-/-^* (*Rag1^-/-^*) mice. OT-I/B6.SJL mice were obtained by crossing OT-I (*Rag1^-/-^*) mice with B6.SJL (CD45.1) mice. Mice were housed in a pathogen-free environment and treated in accordance with the Canadian Council on Animal Care guidelines.

### Cell lines

B16-OVA cells were kindly provided by A. Lamarre (INRS-Institut Armand-Frappier). B16-OVA cells were cultured in DMEM (Corning; REF: 10-017-CV) supplemented with 10% of *Nu* serum (Corning; ref # 355104), 1mM of sodium pyruvate (Corning; REF: 25-000-Cl), penicillin/streptomycin (Corning; REF: 30-002-Cl) in presence of 5 mg/ml of G418 (Corning; CAS#108321-42-2) to select for OVA expressing cells.

### *In vitro* generation of OT-I CD8^+^ effector T cells for anti-tumor ACT

*Nr4a3^+/+^* or *Nr4a3^-/-^* OT-I T cells (CD45.2^+^) were isolated from lymph nodes and mixed with B6.SJL splenocytes (CD45.1^+^) cells in 4:6 ratio. Stimulation was performed on 6-well plate at 5 x 10^6^ cells per ml in the presence of 0.1 µg/ml of OVA peptide (Midwest Biotech) in complete RPMI media (Corning; ref # 10-040-CV) that is a RPMI supplemented with 10% of *Nu* serum (Corning; ref # 355104), 1mM of sodium pyruvate (Corning; ref # 25-000-Cl), 10mM of HEPES (Corning; ref # 25-060-Cl), 0,1 mM of MEM non-essential amino acids (Gibco Life Technologies; ref # 11140-050), 2mM of glutamine (Corning; ref # 25-005-Cl), 10µM of 2-mercaptoethanol (Gibco Life Technologies; ref # 21985-023) and penicillin/streptomycin (Corning; ref # 30-002-Cl). 24h post stimulation, culture media was removed, and the stimulated cells were gently washed without disrupting their adherence to the plate. New media was added, and the culture continued for an additional 24h. At 48h post stimulation cells were harvested, washed, and resuspended at 1 x 10^6^ cells/ml. Recombinant rhIL-2 (Novartis; PROLEUKIN®) was added to the cell suspension at 100U/ml and the culture was continued for an additional day before the adoptive transfer.

### ACT and anti-PD-L1 treatment

10^6^ *in vitro Nr4a3^+/+^* or *Nr4a3^-/-^* OT-I effectors were adoptively transferred i.v. into B6.SJL mice injected sub-cutaneously with 5 x 10^5^ B16-OVA cells into the right flank seven days before. The anti-PD-L1 treatment was then added by injecting the anti-PD-L1 antibody (BioXcell; clone 10F.9G2) or its isotype (BioXcell or Leinco; Rat IgG2b) control at day 7, 10, 13, 16 post-tumor implantation by intraperitoneal injection of 200 µg of antibody per mouse. The tumor measurement was performed by measuring of the 2-perpendicular axis and the area was calculated by multiplying the 2 axes^8^. For survival curve experiments, the tumor growth was followed every 2 days until the experimental endpoint (200 mm^2^ or tumor ulceration). For TILs phenotyping experiments, at day 12-post-tumor implementation, tumor-bearing mice were treated with 10^6^ *in vitro* generated *Nr4a3^+/+^*or *Nr4a3^-/-^* OT-I effectors. These mice were then treated with anti-PD-L1 at day 15 and 18 post-tumor implantation (200 µg i.p. per mouse) and the tumors were harvested at day 21 post-tumor injection for TIL characterization. For the TIL phenotyping experiments tumors with an area lower than 30mm^2^ were excluded from the analysis to compare the immune response to a well-established tumor burden.

### TILs preparation for flow cytometry analysis

Tumors were extracted, disrupted (between 2 frosted glass slides) and digested with 1mg/ml of collagenase D (Sigma Life Science; ref # 11088882001) and 100µg/ml of DNAse I (Sigma Life Science; ref # D5025-375KU) for 15 min at 37°C^28^. The tumor cell suspension was filtered (100µm) and red blood cells were lysed using 0.83% NH_4_Cl (Bio Basic; CAS #12125-02-9) for 5 min at RT. Prior to antibody stainings, tumor cell suspensions were incubated 10 min at RT with Fc block (Leico; C381) and a viability dye (Biolegend; Zombie Aqua (ref # 423102) or Zombie NIR (ref # 423106)) to exclude dead cells. Extracellular staining was performed 20 min at 4°C in FACS WASH (FW) buffer as previously described^29^. Transcription factors intranuclear staining was performed using *Foxp3/Transcription Factor Staining Buffer Set* (Invitrogen by Thermo Fisher Scientific; ref # 00-5523-00) according to manufacturer instructions. Flow cytometry analysis was performed on BD LSR II, BD LSRFortessa X-20 and HTS BD LSRFortessa X-20 from BD Biosciences. Data were analyzed using FlowJo software (Tree Star). List of antibodies and flow cytometry reagents is provided in the supplementary materials.

### TILs restimulation for cytokine production analysis

Tumor suspensions were stimulated with 50ng/ml of phorbol 12-myristate13-acetate (Sigma; ref # P8139) and 500ng/ml of ionomycin (Sigma; ref # I0634) in the presence of 10 µg/ml of brefeldin A (Fisher; ref # AAJ62340MB) for 5h at 37°C and 5% CO_2_. Cells were fixed 20 min with 2% paraformaldehyde at RT. Fixed cells were permeabilized with 0.5 % of saponin (Sigma Life Science; ref # S-7900) in FW for 10 min at RT. Permeabilized cells were stained with anti-cytokine antibodies followed by cell surface staining as previously described^29,30^.

### Cell sorting

OT-I *Nr4a3^+/+^* or *Nr4a3^-/-^* TILs were sorted from freshly prepared day 21 tumor cells suspensions obtained from mice that were treated or not with anti-PDL-1 antibodies. Briefly, tumor cell suspensions were incubated with viability dye and Fc-block antibodies (Leico; C381) for 10 min at RT. Tumor cells suspensions were then stained with extracellular antibodies 20 min at 4°C in sorting buffer (PBS, 1% *Nu* serum, 1mM EDTA, 25mM HEPES). Viable OT-I TILs (Zombie-dye^neg^CD8^+^ CD45.1^-^CD45.2^+^) were sorted in complete RPMI media.

### sc-RNAseq sample preparation and library generation

Tumors from mice that were treated with *in vitro* generated *Nr4a3^+/+^*and *Nr4a3^-/-^* OT-I effectors at day 12 and followed by treatment with anti-PD-L1 or isotype control antibodies on day 15 and 18 were collected at day 21. TIL extraction were performed on each collected tumor and then three tumors from each experimental condition were pooled. Tumor cell suspensions were stained with viability dye and Abs as described above. Viable TILs (ZombieAqua^-^ CD8^+^ CD45.1^-^ CD45.2^+^) were sorted in complete RPMI media and kept on ice. Sorted TILs were then washed in sorting buffer and then blocked with Fc-block for 10 min at 4°C before TotalSeq-HashTag Ab staining 30 min on ice. *Nr4a3^+/+^* were stained with HashTag2 (GGTCGAGAGCATTCA) and *Nr4a3^-/-^* with HashTag3 (CTTGCCGCATGTCAT). After the HashTag staining, TILs were washed once with sorting buffer and twice with 0.04 % of unacetylated BSA containing PBS. TILs were then counted and mixed in a 1:1 ratio (*Nr4a3^+/+^* with *Nr4a3^-/-^*OT-I TILs from mice treated with the isotype control; and *Nr4a3^+/+^* with *Nr4a3^-/-^* OT-I TILs from mice treated with anti-PD-L1). Each TILs mix were diluted at 1 x 10^6^ cells /ml and about 5000 cells of each TILs mix were loaded into the 10x Chromium controller for the individual cell sequencing. For each cell mix a scRNA-seq libraries was generated using the Chromium Next GEM Single Cell 3’ Kit v3.1 (with dual index and 3’ Feature barcoding technology). All library preparation steps were performed according to the manufacturer’s instructions. Each cell was sequenced with 50000 reads per cell and 1000 reads for the HashTag. We obtained the individual transcriptome of a total of 10830 OT-I TILs (3148 and 2723 *Nr4a3^+/+^* cells treated or not with anti-PDL-1; 2215 and 2744 *Nr4a3^-/-^*cells treated or not with anti-PD-L1).

### Single cell RNAseq data analysis

Cellranger-6.1.1’s multi pipeline was used to analyze 10x 3’ Cell Multiplexing data of *Nr4a3^+/+^* and *Nr4a3^-/-^* reads from both isotype control and anti-PD-L1 conditions. The resulting gene expression matrices for each sample were then imported in R package Seurat (v4.1) for quality control and downstream analyses. Each sample was first examined individually to identify and filter out empty/dead cells or doublets based on the number of transcripts and number of unique genes present in each cell and on the percentage of mitochondrial reads. All 4 samples were then merged, and the percentage of mitochondrial reads regressed out. Data was log normalized and highly variable genes identified. The expression level of highly variable genes in the cells was scaled and centered along each gene, and linear dimensionality reduction performed by principal component analysis (PCA). Only the first 20 PCs were selected for subsequent analyses. Clusters were identified using Louvain’s algorithm with a resolution of 0.8, yielding a total of 15 clusters. Differential expression analysis on the resulting clusters was conducted using the Wilcoxon rank-sum test for genes expressed at least in 10% cells within the cluster and with a fold change more than 0.10 (log scale). P-values were adjusted for multiple testing using the Bonferroni correction. Uniform Manifold Approximation and Projection (UMAP) was applied for visualization purposes. DEGs were calculated using the functions FindAllMarkers or FindMarkers (Seurat) for two experimental condition or clusters pairwise comparisons using the adjusted *P value* of less than 0.05 as significant and a log2 fold-change threshold of 0.584963 (FC>1.5) as an enriched gene expression value. Gene-set enrichments were calculated using the *AddModuleScore* function (Seurat). *Gene Ontology* analysis of the upregulated genes was done with *G:Profiler* (https://biit.cs.ut.ee/gprofiler/). Gene set enrichment analysis was performed using *Fast Gene Set Enrichment Analysis* (fgsea 1.20.1) package*. Pseudotime* analysis was done using Monocle.

### Statistical analyses

For multiple groups comparisons one way ANOVA (Kruskal-Wallis) analysis with Dunn’s multiple comparisons was performed. For survival curve comparison a log-rank Mantel–Cox test was used. Paired samples (OT-I and OT-I/B6.SJL comparisons) were analyzed using a paired nonparametric paired T test. Unpaired two groups comparison (with a low number of samples) was performed using a Mann Whitney test. Statistical analyses were performed using Prism software (GraphPad Software). Data were presented as individual samples with mean where each dot represent one mouse on scatter plots and also each line represent one mouse on tumor growth follow-up curves. P values of \**P* < 0.05, \*\**P* < 0.01, \*\*\**P* < 0.001, \*\*\*\**P* < 0.0001 were considered statistically significant.

## Acknowledgements

We thank all laboratory members for helpful discussion. We thank Dr O. Conneely for providing *Nr4a3^-/-^* mice and M.-È. Lebel for advice on the B16-OVA tumor model. We thank Frédéric Duval for cell sorting and the animal care technicians for mice husbandry. We are grateful to S. Boissel, V. Calderon, C. Grou from IRCM for help with the scRNA-seq experiment and bioinformatical analysis.

## Author contribution

L.O., S.B., N.L. designed and analysed the experiments. L.O., D.M.D.S., S.B., S” M. performed the experiments. N.L., L.O. wrote the paper.

## Data and materials availability

The raw scRNA-seq will be deposited at Gene Expression Omnibus.

---

**Figure. S1.**
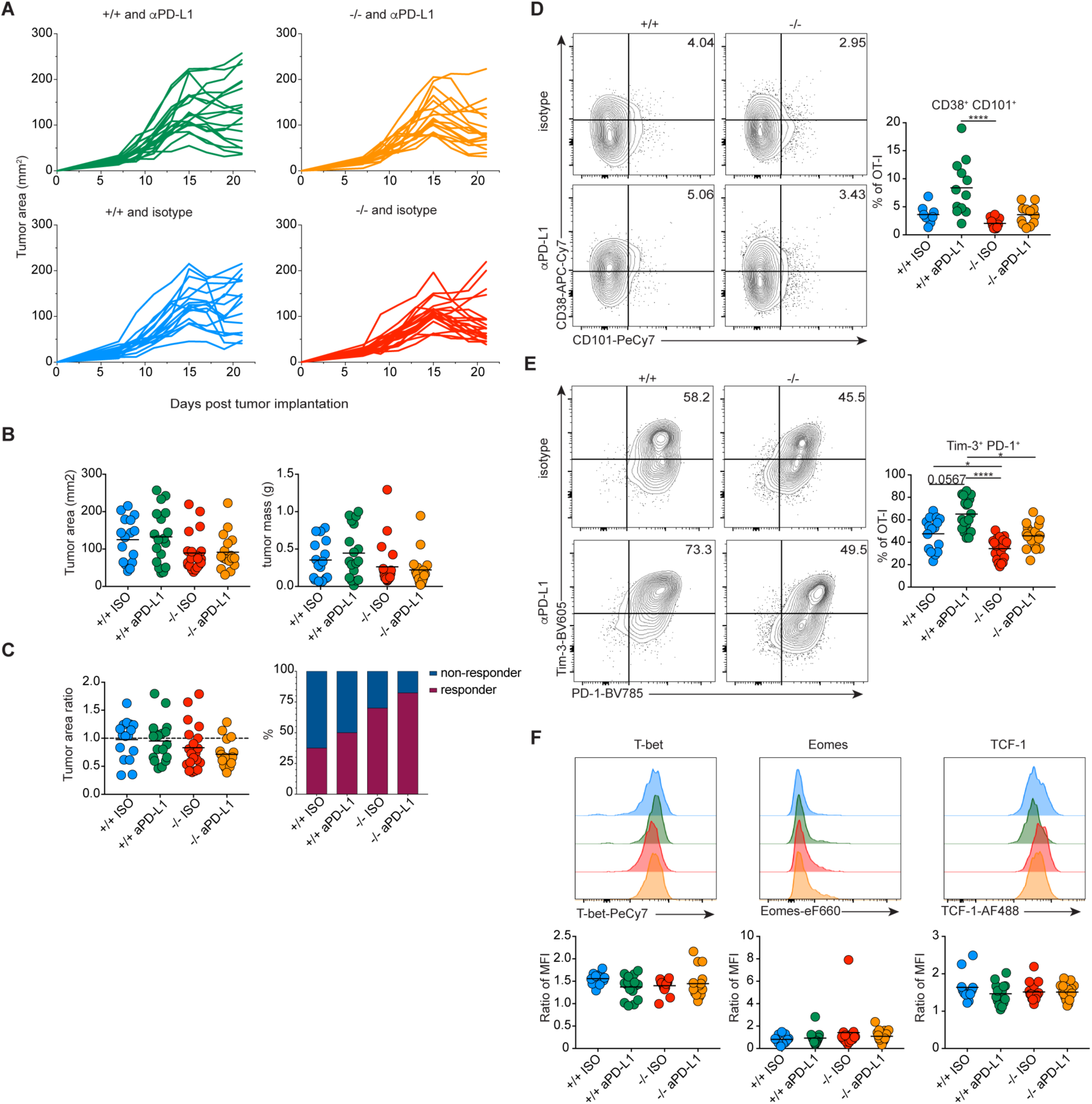
ACT with NR4A3-deficient CD8^+^ T cells increases tumor control and decreases T cell exhaustion. **A.** Tumor growth curves of mice implanted with B16-OVA melanoma cells and treated 12 days later with *Nr4a3*^+/+^ or *Nr4a3*^-/-^ *in vitro* generated OT-I effectors and cotreated on day 15 and 18 with anti-PD-L1 or isotype control (ISO). **B.** Tumor area and tumor mass of tumors extracted on day 21. **C**. The proportion of mice responding to the therapy was evaluated by the ratio of the tumor area on day 21 compared to the tumor area on day 15. Co-expression of CD38 and CD101 (**D**) or PD-1 and Tim-3 (**E**) by *Nr4a3*^+/+^ or *Nr4a3*^-/-^ OT-I TILs at day 21. **F.** T-bet, Eomes, and TCF-1 expression by *Nr4a3*^+/+^ or *Nr4a3*^-/-^ TILs at day 21. Data are from 3 to 4 independent experiments. Each dot represents one mouse. In the tumor growth curves, each line represents one mouse. Kruskal-Wallis ANOVA with Dunn’s multiple comparisons was used for multiple groups comparison: \**P*<0.05, \*\*\*\**P*<0.0001.

**Figure. S2.**
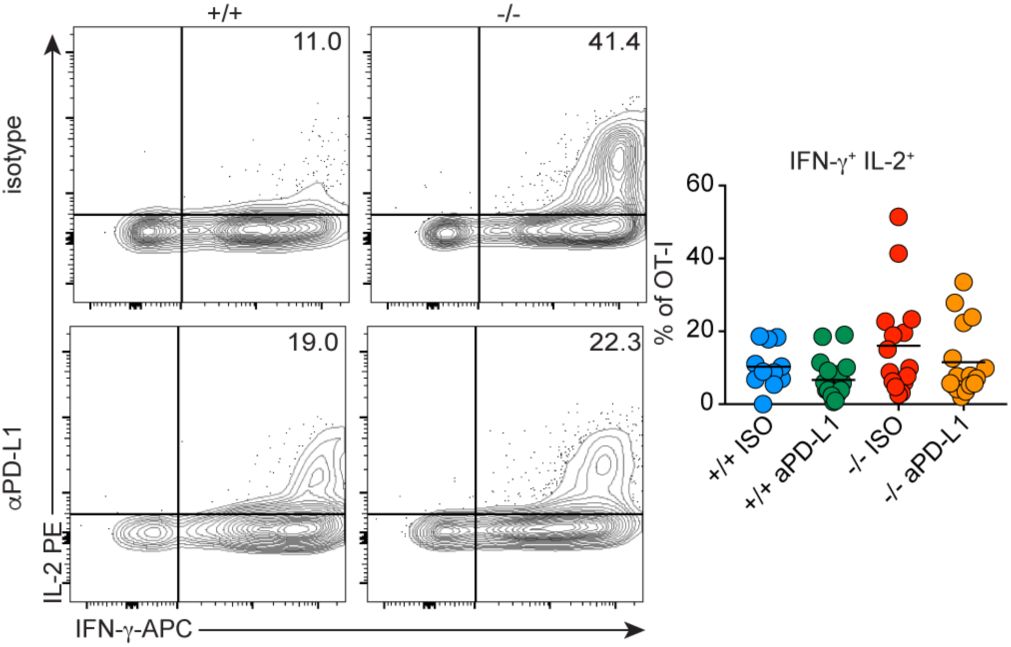
NR4A3 deficiency does not impact the proportion of IL-2^+^IFN-γ^+^ TILs. Mice were implanted with B16-OVA melanoma cells and treated 12 days later with *Nr4a3*^+/+^ or *Nr4a3*^-/-^ *in vitro* generated OT-I effectors followed by anti-PD-L1 or isotype control treatment on day 15 and 18. At day 21, tumor cell suspensions were restimulated for 5h with PMA/ionomycin for cytokine production quantification. Proportions of IL-2^+^IFN-γ^+^ producing OT-I TILs. Each dot represents one mouse. Kruskal-Wallis ANOVA with Dunn’s multiple comparisons was used for multiple groups comparison.

**Figure. S3.**
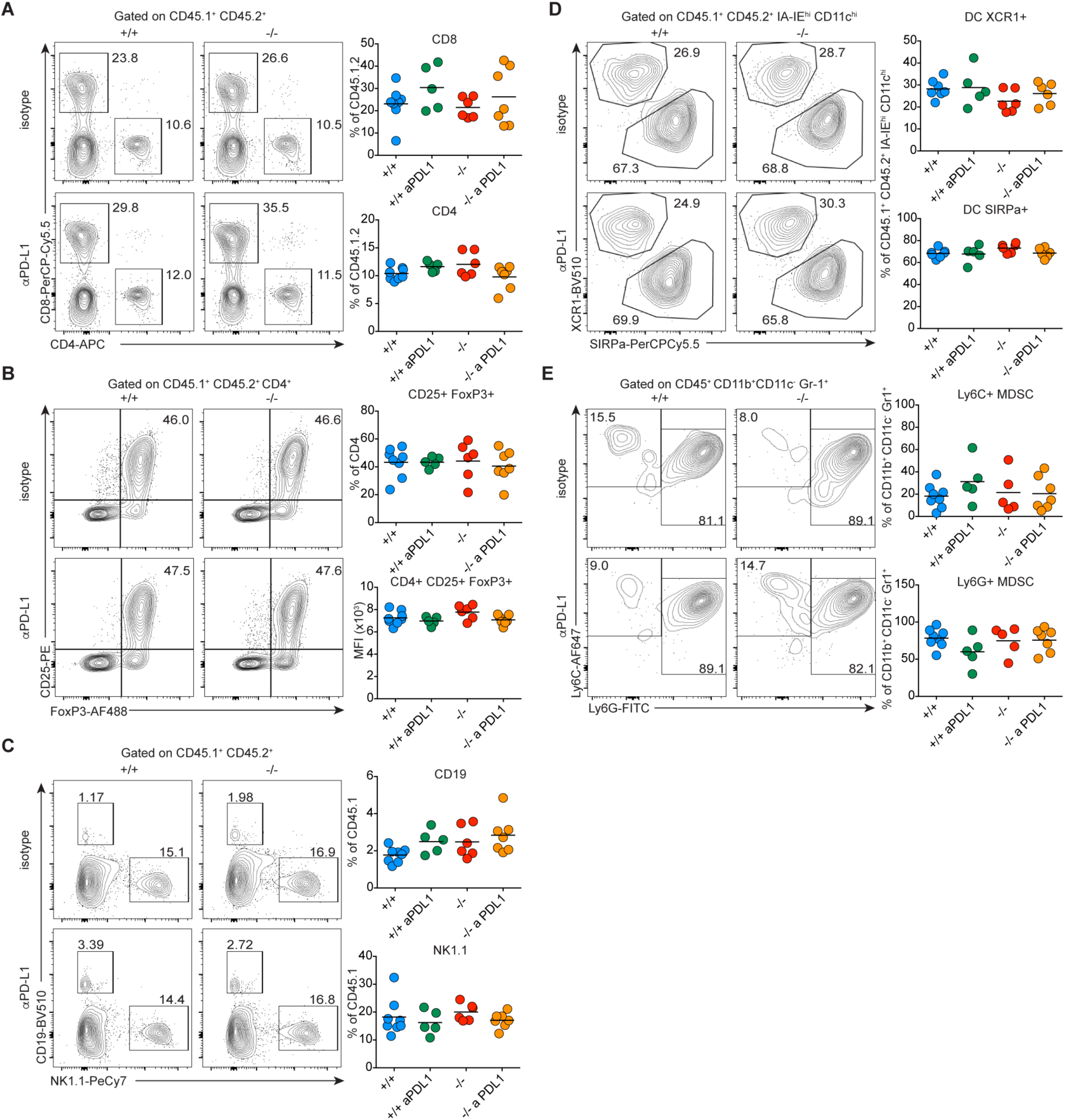
NR4A3 deficiency does not affect tumor infiltration by other immune cell types. At day 21 post-tumor implantation, the proportions of endogenous tumor infiltrating of CD8^+^ and CD4^+^ T cells (**A**), regulatory T cells (**B**), B and NK cells (**C**), dendritic cells (DCs) (**D**), and myeloid-derived suppressor cells (MDSCs) (**E**) were analyzed by cytometry. Data are from 1 experiment, and each dot represents one mouse. Kruskal-Wallis ANOVA with Dunn’s multiple comparisons was used for multiple groups comparison.

**Figure. S4.**
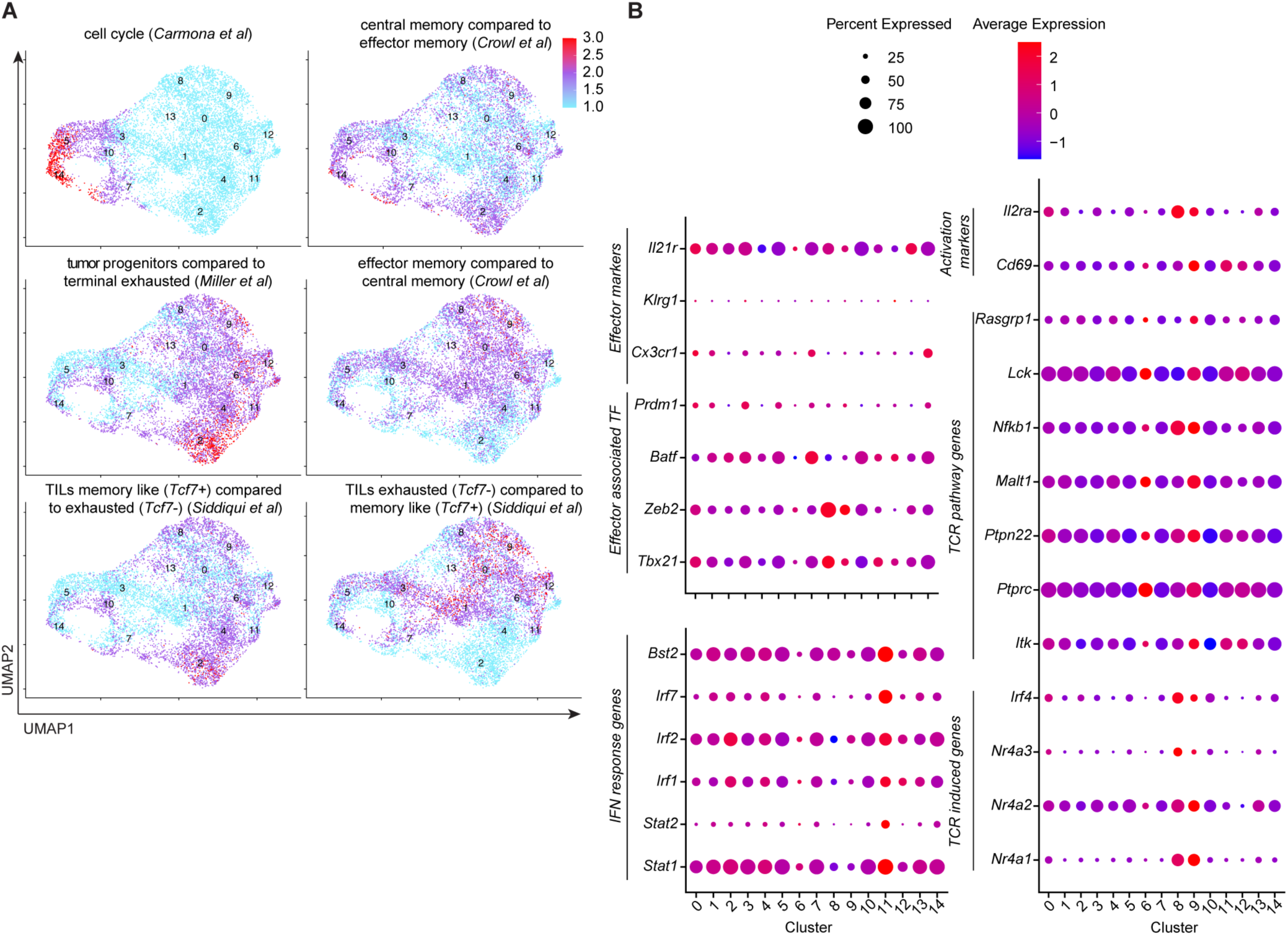
*Nr4a3*^+/+^ or *Nr4a3*^-/-^ TILs transcriptional profile under anti-PD-L1 treatment. Mice were implanted with B16-OVA melanoma cells and treated 12 days later with *Nr4a3*^+/+^ or *Nr4a3*^-/-^ *in vitro* generated OT-I effectors, as well as with anti-PD-L1 or isotype control on day 15 and 18. At day 21, OT-I TILs were isolated for scRNA-seq analysis. **A**. Gene signature projection of different CD8^+^ T cell subsets on the scRNA-seq UMAP. **B.** Dot plot quantification of selected gene expression by the different scRNA-seq clusters. Data are from one scRNA-seq experiment where 3 independent biological samples were pooled for each treatment condition.

**Figure. S5.**
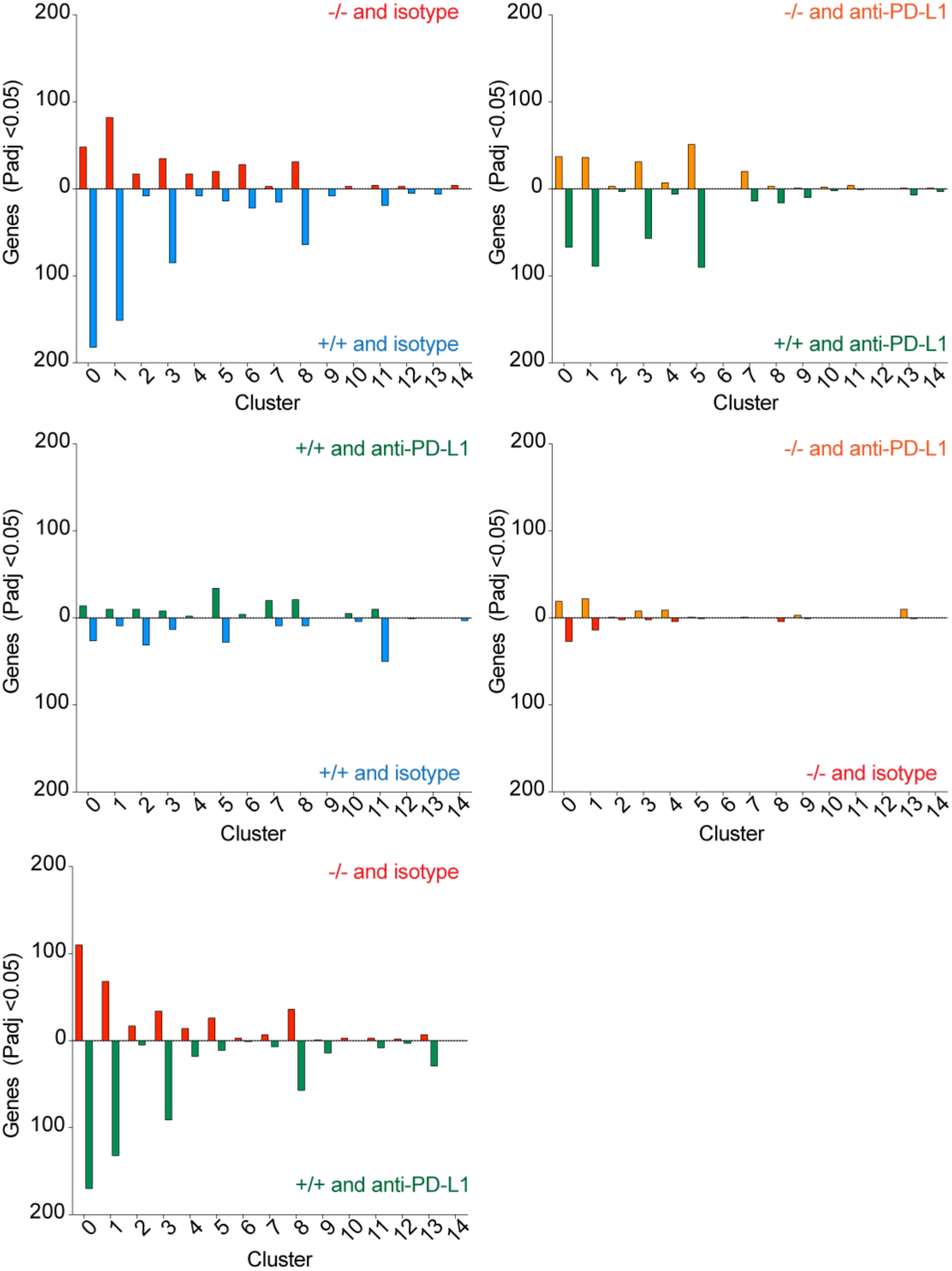
*Nr4a3*^+/+^ or *Nr4a3*^-/-^ TILs differentially expressed genes by clusters. Mice were implanted with B16-OVA melanoma cells and 12 days later treated with *Nr4a3*^+/+^ or *Nr4a3*^-/-^ *in vitro* generated OT-I effectors, as well as with anti-PD-L1 or isotype control on day 15 and 18. At day 21, OT-I TILs were isolated for scRNA-seq analysis. Numbers of differentially expressed genes (*Padj*<0.05) in each cluster in function of the type of treatment (genotype and PD-L1 blockade). Data are from one scRNA-seq experiment where 3 independent biological samples were pooled for each treatment condition.

**Figure. S6.**
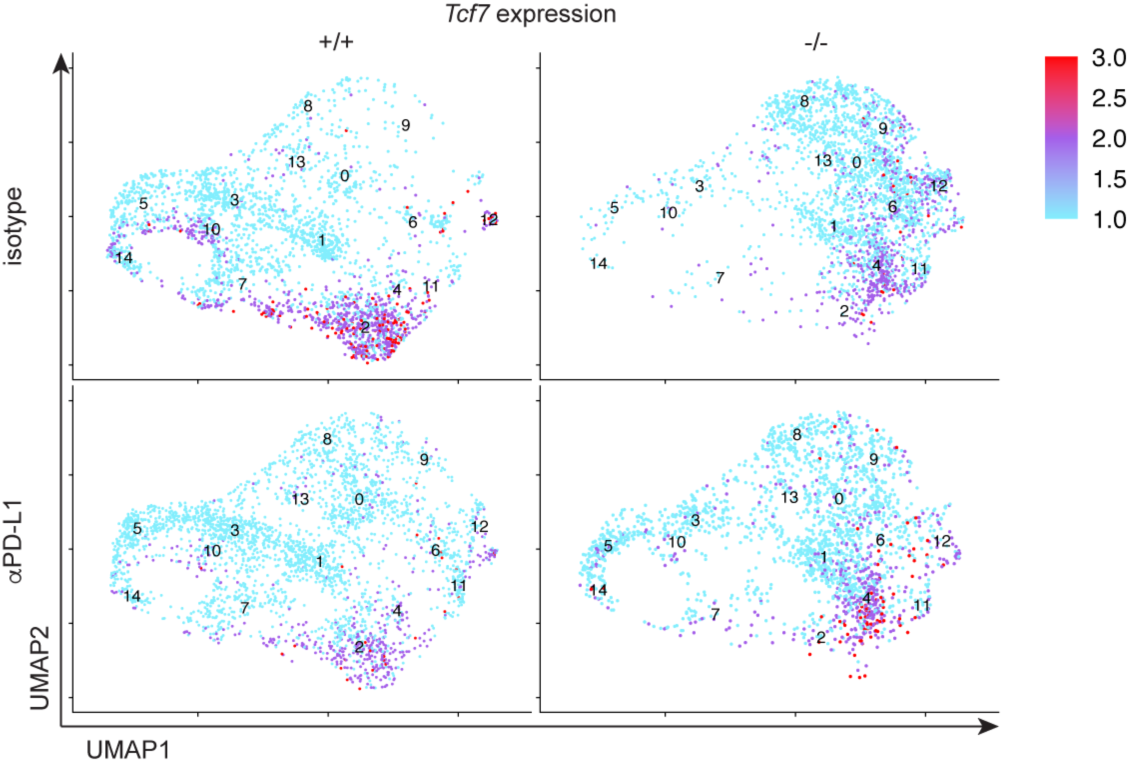
*Tcf7* expression by *Nr4a3*^+/+^ or *Nr4a3*^-/-^ OT-I TILs. *Tcf7* gene expression projection on the UMAP of *Nr4a3*^+/+^ or *Nr4a3*^-/-^ TILs under anti-PD-L1 or isotype control treatment. Data are from one scRNA-seq experiment where 3 independent biological samples were pooled for each treatment condition.

**Table SI.**
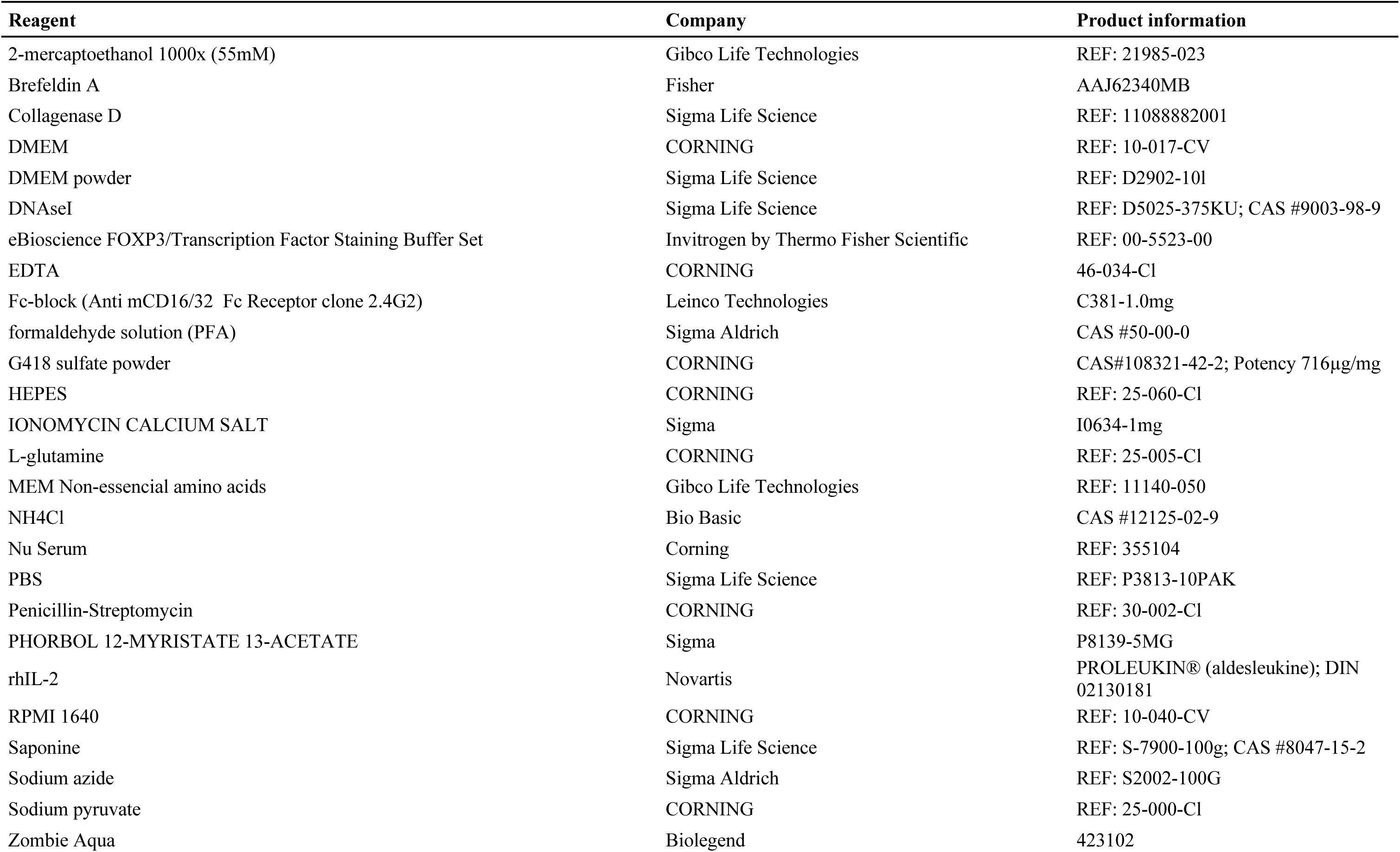
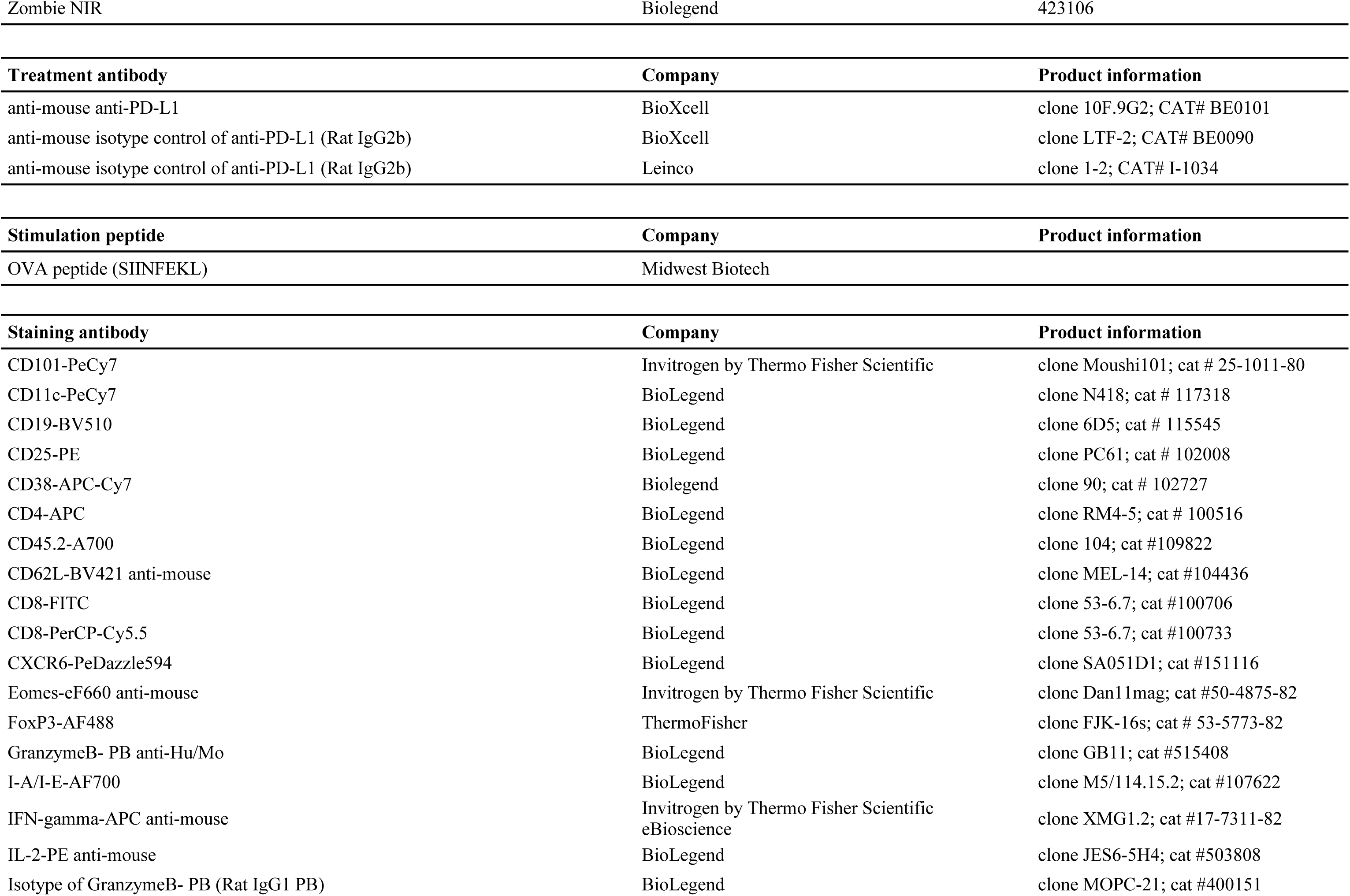
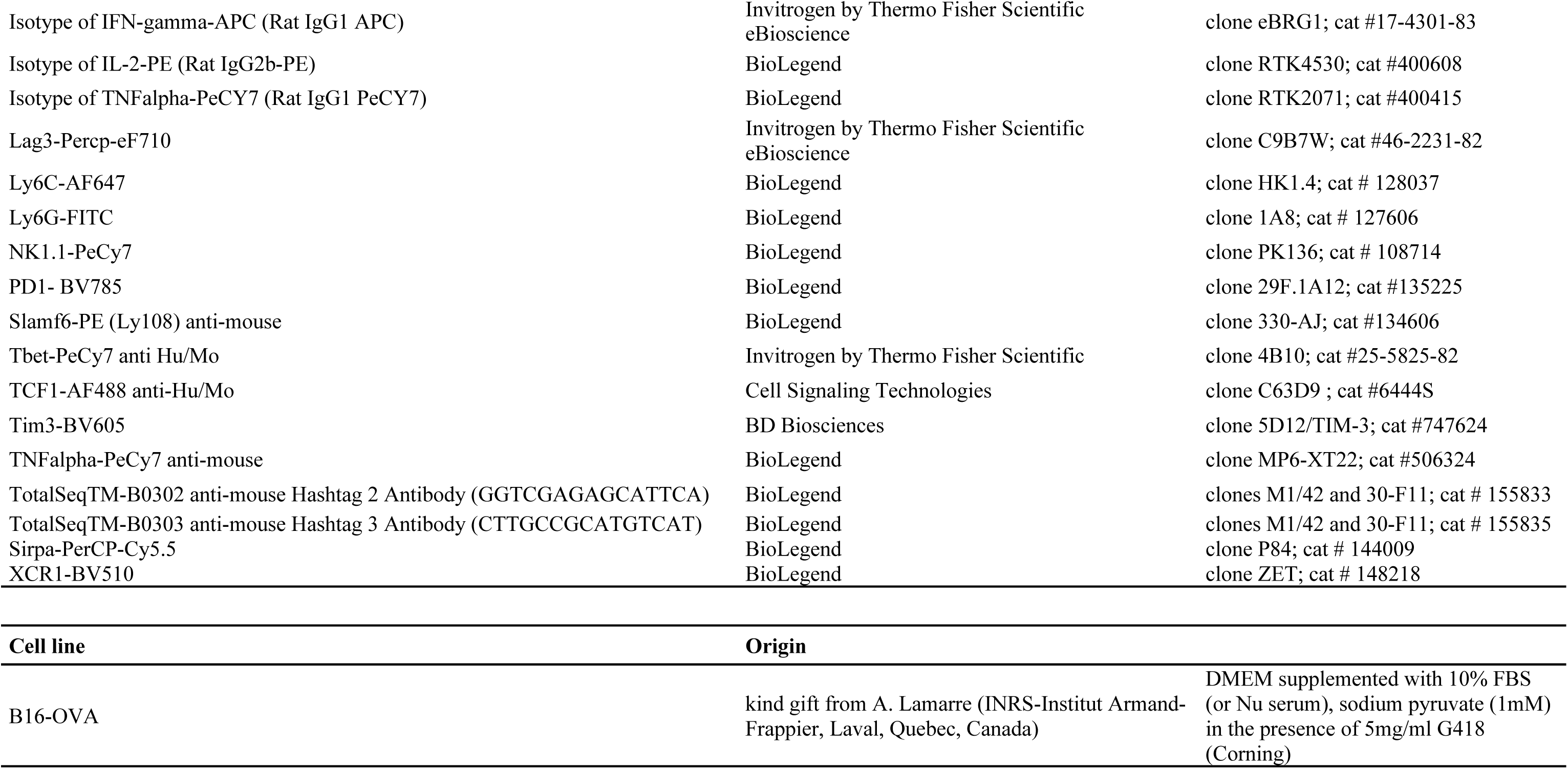
Antibodies and reagents list.

