## Supplemental Table for "NR4A3 deficiency in CD8^+^ T cells improves adoptive T cell therapy of cancer"

**SUPPLEMENTARY TABLE**

Table SI. Antibodies and reagents list

**Table SI. Antibodies and reagents list**

| **Reagent** | **Company** | **Product information** |
| --- | --- | --- |
| 2-mercaptoethanol 1000x (55mM) | Gibco Life Technologies | REF: 21985-023 |
| Brefeldin A | Fisher | AAJ62340MB |
| Collagenase D | Sigma Life Science | REF: 11088882001 |
| DMEM | CORNING | REF: 10-017-CV |
| DMEM powder | Sigma Life Science | REF: D2902-10l |
| DNAseI | Sigma Life Science | REF: D5025-375KU; CAS #9003-98-9 |
| eBioscience FOXP3/Transcription Factor Staining Buffer Set | Invitrogen by Thermo Fisher Scientific | REF: 00-5523-00 |
| EDTA | CORNING | 46-034-Cl |
| Fc-block (Anti mCD16/32 Fc Receptor clone 2.4G2) | Leinco Technologies | C381-1.0mg |
| formaldehyde solution (PFA) | Sigma Aldrich | CAS #50-00-0 |
| G418 sulfate powder | CORNING | CAS#108321-42-2; Potency 716µg/mg |
| HEPES | CORNING | REF: 25-060-Cl |
| IONOMYCIN CALCIUM SALT | Sigma | I0634-1mg |
| L-glutamine | CORNING | REF: 25-005-Cl |
| MEM Non-essencial amino acids | Gibco Life Technologies | REF: 11140-050 |
| NH4Cl | Bio Basic | CAS #12125-02-9 |
| Nu Serum | Corning | REF: 355104 |
| PBS | Sigma Life Science | REF: P3813-10PAK |
| Penicillin-Streptomycin | CORNING | REF: 30-002-Cl |
| PHORBOL 12-MYRISTATE 13-ACETATE | Sigma | P8139-5MG |
| rhIL-2 | Novartis | PROLEUKIN® (aldesleukine); DIN 02130181 |
| RPMI 1640 | CORNING | REF: 10-040-CV |
| Saponine | Sigma Life Science | REF: S-7900-100g; CAS #8047-15-2 |
| Sodium azide | Sigma Aldrich | REF: S2002-100G |
| Sodium pyruvate | CORNING | REF: 25-000-Cl |
| Zombie Aqua | Biolegend | 423102 |
| Zombie NIR | Biolegend | 423106 |
| **Treatment antibody** | **Company** | **Product information** |
| anti-mouse anti-PD-L1 | BioXcell | clone 10F.9G2; CAT# BE0101 |
| anti-mouse isotype control of anti-PD-L1 (Rat IgG2b) | BioXcell | clone LTF-2; CAT# BE0090 |
| anti-mouse isotype control of anti-PD-L1 (Rat IgG2b) | Leinco | clone 1-2; CAT# I-1034 |
| **Stimulation peptide** | **Company** | **Product information** |
| OVA peptide (SIINFEKL) | Midwest Biotech |  |
| **Staining antibody** | **Company** | **Product information** |
| CD101-PeCy7 | Invitrogen by Thermo Fisher Scientific | clone Moushi101; cat # 25-1011-80 |
| CD11c-PeCy7 | BioLegend | clone N418; cat # 117318 |
| CD19-BV510 | BioLegend | clone 6D5; cat # 115545 |
| CD25-PE | BioLegend | clone PC61; cat # 102008 |
| CD38-APC-Cy7 | Biolegend | clone 90; cat # 102727 |
| CD4-APC | BioLegend | clone RM4-5; cat # 100516 |
| CD45.2-A700 | BioLegend | clone 104; cat #109822 |
| CD62L-BV421 anti-mouse | BioLegend | clone MEL-14; cat #104436 |
| CD8-FITC | BioLegend | clone 53-6.7; cat #100706 |
| CD8-PerCP-Cy5.5 | BioLegend | clone 53-6.7; cat #100733 |
| CXCR6-PeDazzle594 | BioLegend | clone SA051D1; cat #151116 |
| Eomes-eF660 anti-mouse | Invitrogen by Thermo Fisher Scientific | clone Dan11mag; cat #50-4875-82 |
| FoxP3-AF488 | ThermoFisher | clone FJK-16s; cat # 53-5773-82 |
| GranzymeB- PB anti-Hu/Mo | BioLegend | clone GB11; cat #515408 |
| I-A/I-E-AF700 | BioLegend | clone M5/114.15.2; cat #107622 |
| IFN-gamma-APC anti-mouse | Invitrogen by Thermo Fisher Scientific eBioscience | clone XMG1.2; cat #17-7311-82 |
| IL-2-PE anti-mouse | BioLegend | clone JES6-5H4; cat #503808 |
| Isotype of GranzymeB- PB (Rat IgG1 PB) | BioLegend | clone MOPC-21; cat #400151 |
| Isotype of IFN-gamma-APC (Rat IgG1 APC) | Invitrogen by Thermo Fisher Scientific eBioscience | clone eBRG1; cat #17-4301-83 |
| Isotype of IL-2-PE (Rat IgG2b-PE) | BioLegend | clone RTK4530; cat #400608 |
| Isotype of TNFalpha-PeCY7 (Rat IgG1 PeCY7) | BioLegend | clone RTK2071; cat #400415 |
| Lag3-Percp-eF710 | Invitrogen by Thermo Fisher Scientific eBioscience | clone C9B7W; cat #46-2231-82 |
| Ly6C-AF647 | BioLegend | clone HK1.4; cat # 128037 |
| Ly6G-FITC | BioLegend | clone 1A8; cat # 127606 |
| NK1.1-PeCy7 | BioLegend | clone PK136; cat # 108714 |
| PD1- BV785 | BioLegend | clone 29F.1A12; cat #135225 |
| Slamf6-PE (Ly108) anti-mouse | BioLegend | clone 330-AJ; cat #134606 |
| Tbet-PeCy7 anti Hu/Mo | Invitrogen by Thermo Fisher Scientific | clone 4B10; cat #25-5825-82 |
| TCF1-AF488 anti-Hu/Mo | Cell Signaling Technologies | clone C63D9 ; cat #6444S |
| Tim3-BV605 | BD Biosciences | clone 5D12/TIM-3; cat #747624 |
| TNFalpha-PeCy7 anti-mouse | BioLegend | clone MP6-XT22; cat #506324 |
| TotalSeqTM-B0302 anti-mouse Hashtag 2 Antibody (GGTCGAGAGCATTCA) | BioLegend | clones M1/42 and 30-F11; cat # 155833 |
| TotalSeqTM-B0303 anti-mouse Hashtag 3 Antibody (CTTGCCGCATGTCAT)  Sirpa-PerCP-Cy5.5  XCR1-BV510 | BioLegend  BioLegend  BioLegend | clones M1/42 and 30-F11; cat # 155835  clone P84; cat # 144009  clone ZET; cat # 148218 |
| **Cell line** | **Origin** |  |
| B16-OVA | kind gift from A. Lamarre (INRS-Institut Armand-Frappier, Laval, Quebec, Canada) | DMEM supplemented with 10% FBS (or Nu serum), sodium pyruvate (1mM) in the presence of 5mg/ml G418 (Corning) |
